# Inhibition of SARS-CoV-2 polymerase by nucleotide analogs: a single molecule perspective

**DOI:** 10.1101/2020.08.06.240325

**Authors:** Mona Seifert, Subhas Chandra Bera, Pauline van Nies, Robert N. Kirchdoerfer, Ashleigh Shannon, Thi-Tuyet-Nhung Le, Xiangzhi Meng, Hongjie Xia, James M. Wood, Lawrence D. Harris, Flávia S. Papini, Jamie J. Arnold, Steven C. Almo, Tyler L. Grove, Pei-Yong Shi, Yan Xiang, Bruno Canard, Martin Depken, Craig E. Cameron, David Dulin

## Abstract

The nucleotide analog Remdesivir (RDV) is the only FDA-approved antiviral therapy to treat infection by severe acute respiratory syndrome coronavirus 2 (SARS-CoV-2). The physical basis for efficient utilization of RDV by SARS-CoV-2 polymerase is unknown. Here, we characterize the impact of RDV and other nucleotide analogs on RNA synthesis by the polymerase using a high-throughput, single-molecule, magnetic-tweezers platform. The location of the modification in the ribose or in the base dictates the catalytic pathway(s) used for its incorporation. We reveal that RDV incorporation does not terminate viral RNA synthesis, but leads the polymerase into deep backtrack, which may appear as termination in traditional ensemble assays. SARS-CoV-2 is able to evade the endogenously synthesized product of the viperin antiviral protein, ddhCTP, though the polymerase incorporates this nucleotide analog well. This experimental paradigm is essential to the discovery and development of therapeutics targeting viral polymerases.

**Teaser:** We revise Remdesivir’s mechanism of action and reveal SARS-CoV-2 ability to evade interferon-induced antiviral ddhCTP

## Introduction

SARS-CoV-2 has infected more than 100 million humans worldwide, causing over 2 million deaths, with numbers still on the rise. We are currently living through the third coronavirus outbreak in less than twenty years, and we are desperately in need of broad-spectrum antiviral drugs that are capable of targeting this emerging family of human pathogens. To this end, nucleotide analogs (NAs) represent a powerful approach, as they target the functionally and structurally conserved coronavirus polymerase, and their insertion in the viral RNA induces either premature termination or a lethal increase in mutations. The coronavirus polymerase is composed of the nsp12 RNA-dependent RNA polymerase (RdRp), and the nsp7 and nsp8 co-factors, with a stoichiometry of 1:1:2 (*1-4*). This polymerase is thought to associate with several additional viral proteins, including the nsp13, a 5’-to-3’ RNA helicase (*5, 6*) and the nsp14, a 3’-to-5’ exoribonuclease (*7-12*). The latter proofreads the RNA synthesized by the polymerase and associated factors (*13*), a unique feature of coronaviruses relative to all other families of RNA viruses. Proofreading likely contributes to the stability of the unusually large, ∼30 kb, coronavirus genome. In addition, proofreading may elevate the tolerance of coronaviruses to certain NAs (e.g., ribavirin (*14*)) and therefore should be considered in the development of potent NAs. Remdesivir (RDV) is a recently discovered NA that showed efficacy against Ebola infection (*15*) and has been successfully repurposed for the treatment of SARS-CoV-2 infection (*7, 16-20*). The success of RDV relies on its efficient incorporation by the polymerase (*17, 21*) and probable evasion of the proofreading machinery (*7*). Understanding how RDV achieves these two tasks, will help to guide the rational design of more efficacious NAs for the current and future outbreaks. To this end, it is essential to build a comprehensive model describing the selection and incorporation mechanisms that control the utilization of NAs by the coronavirus polymerase and to define the determinants of the base and ribose responsible for selectivity and potency. Creation of such a model requires development of an experimental paradigm for the coronavirus polymerase that permits monitoring of hundreds to thousands of nucleotide incorporation events in the presence of all the natural ribonucleotides at physiological concentration. Here, we present such an experimental paradigm that provides insights into the mechanism and efficacy of current and underexplored NAs.

## Results

### Monitoring SARS-CoV-2 RNA synthesis at the single-molecule level

To enable the observation of rare events, such as nucleotide mismatch and NA incorporation, even in the presence of saturating NTP concentration, we have developed a single-molecule, high-throughput, magnetic tweezers assay to monitor SARS-CoV-2 RNA synthesis activity at near single base resolution (*22*). A SARS-CoV-2 polymerase formed of nsp12, nsp7, and nsp8 assembles and initiates RNA synthesis at the 3’-end of the magnetic bead-attached handle, and converts the 1043 nt long single-stranded (ss) RNA template into a double-stranded (ds) RNA in the presence of NTPs (**Fig. 1A, Fig. S1A, Materials and Methods**). The conversion from ssRNA to dsRNA displaces the magnetic bead along the vertical axis and directly informs on the number of incorporated nucleotides (**Fig. 1A, Materials and Methods**) (*23*). During each experiment, hundreds of magnetic beads are followed in parallel (**Fig. S1C**), yielding dozens of traces of SARS-CoV-2 polymerase activity per experiment (**Fig. 1B**). As previously observed for other viral RdRps (*22-24*), the traces reveal substantial, heterogeneous dynamics, with bursts of activity interrupted by pauses of duration varying from ∼0.5 s to ∼60 s in **Fig. 1B**. This dynamic is intrinsic to the polymerase elongation kinetics and does not result from viral proteins exchange (*25*). To extract the elongation kinetics of the SARS-CoV-2 polymerase, we scanned the traces with non-overlapping 10-nt windows to measure the duration of time required to complete the ten successive nucleotide-incorporation events. Each duration of time has been coined a dwell time, which is the kinetic signature of the rate-limiting event of the 10 nt addition, i.e. the 10 nucleotide addition cycles themselves, or a pause (*22-24*) (**Materials and Methods**). We fitted the distribution of dwell times using the stochastic-pausing model that describes well the kinetics of nucleotide addition of the coronavirus polymerase (*25*) and other viral RdRps (*22-24, 26*) (**Fig. 1C, Materials and Methods**). This model is composed of four distributions: a pause-free nucleotide addition rate, Pause 1, Pause 2, and the backtrack pauses (**Fig. 1C, Supplementary Materials**). Statistics and all parameter values extracted from the analysis are reported in **Table S1** and **Table S2**. SARS-CoV-2 polymerase elongation kinetics is robustly described by a model where the nucleotide addition rate is the kinetic signature of the nucleotide addition burst (NAB) pathway, from which the RdRp stochastically and rarely switches into the slow nucleotide addition (SNA) pathway, and even more rarely into the very slow nucleotide addition (VSNA) pathway, the latter being consistent in rate and probability with mismatch incorporation (**Fig. 1D**) (*25*). Pause 1 and Pause 2 are respectively the kinetic signatures of the SNA and VSNA pathways (**Fig. 1D**), while the long-lived pauses relate to a catalytically incompetent polymerase backtrack state (**Supplementary Materials**) (*25*). Increasing the temperature from 25°C to 37°C, SARS-CoV-2 polymerase reveals a strong temperature dependence, which translates into a 2-fold decrease in the median replication time (**Fig. 1E**), while not affecting the RNA synthesis product length (**Fig. 1F**). Analyzing the dwell-time distribution at 25°C and 37 °C (**Fig. 1G**), we extracted a ∼2.6-fold enhancement in nucleotide addition rate, from (65.6 ± 0.5) *nt. s*^−1^ to (169.0 ± 3.8) *nt*. *s*^−1^, making the SARS-CoV-2 polymerase the fastest RNA polymerase characterized to date (**Fig. 1H**) (*13, 21*). Pause 1 and Pause 2 exit rates also increased by ∼3-fold (**Fig. 1H**), whereas their respective probabilities increased by 2- and 5-fold (**Fig. S1D**). The latter results are rather surprising, as poliovirus and human rhinovirus C RdRps showed only an exit rate increase with no change in probability (*24*).

**Fig. 1.**
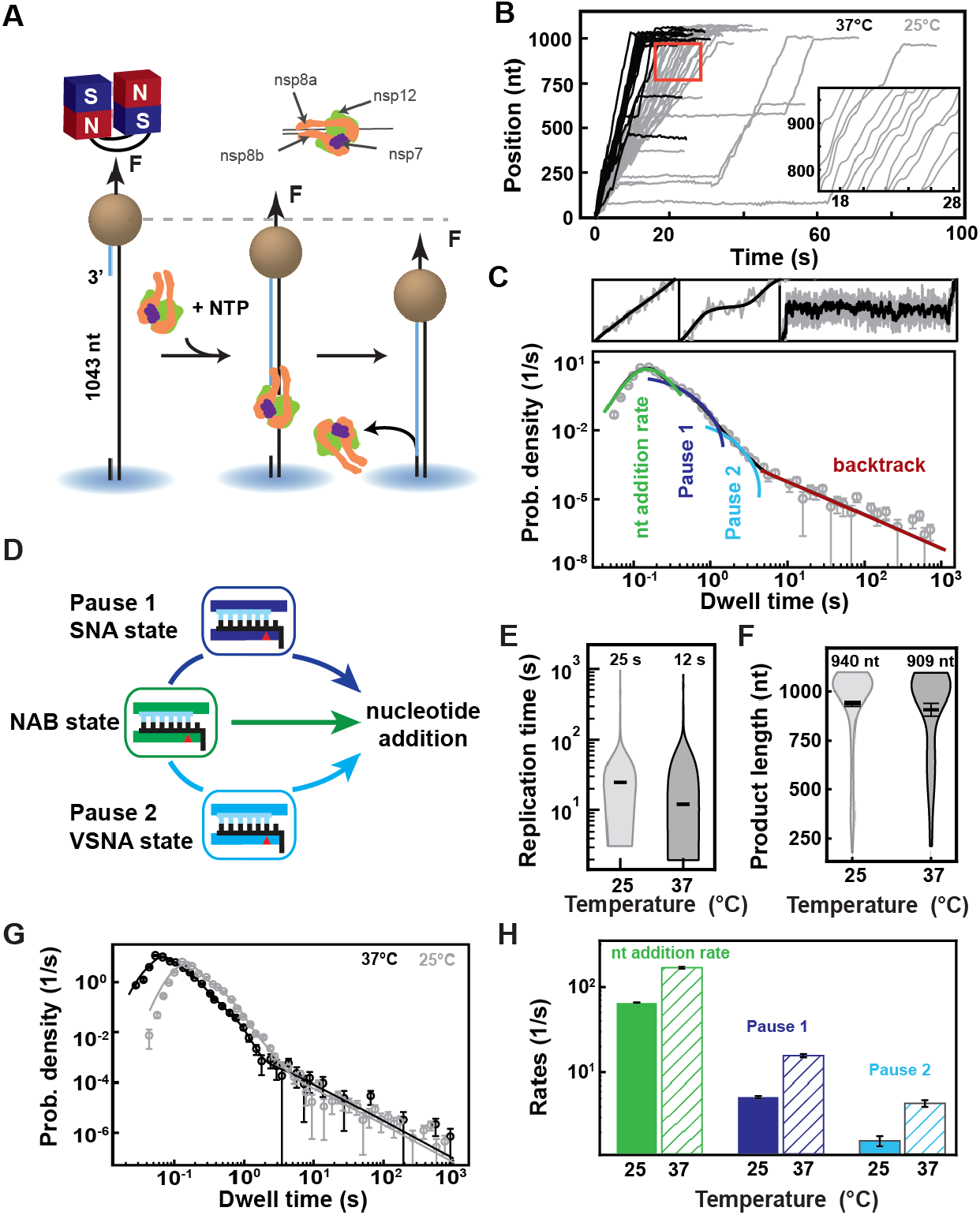
SARS-CoV-2 polymerase is a fast and processive RNA polymerase complex. **(A)** Schematic of the magnetic tweezers assay to monitor RNA synthesis by the SARS-CoV-2 polymerase complex. A magnetic bead is attached to a glass coverslip surface by a 1043 long ssRNA construct that experiences a constant force F. The polymerase, formed by nsp7, nsp8, and nsp12, assembles at the 3’-end of the RNA strand annealed to the template. The subsequent conversion of the ssRNA template into dsRNA reduces the end-to-end extension of the tether, signaling replication activity. **(B)** SARS-CoV-2 polymerase activity traces acquired at either 25°C (gray) or 37°C (black), showing bursts of nucleotide addition interrupted by pauses. The inset is a zoom-in of the traces captured in the red square. **(C)** The dwell times collected from (B) are assembled into a distribution that is fitted using a pause-stochastic model (**Supplementary Materials**) (solid lines). The model includes four different probability distribution functions that describe the event that kinetically dominates the dwell time: uninterrupted ten nucleotide additions (green), exponentially distributed Pause 1 and 2 (blue and cyan, respectively), and the power law distributed backtrack (red). **(D)** The dwell time distribution in (C) is described by the viral RdRp kinetic model (adapted from Ref. (*22*)). Fast nucleotide addition is achieved by the nucleotide addition burst (NAB) pathway with the nucleotide addition rate extracted from (C). Pause 1 and Pause 2 are the kinetic signature of the slow and very slow nucleotide addition (SNA and VSNA, respectivelly) pathways, the latter being likely related to nucleotide mismatch incorporation. **(E)** Total replication time and **(F)** product length of SARS-CoV-2 polymerase activity traces at either 25°C or 37°C. The median total replication time and mean product length are indicated above the violin plots, and represented as thick horizontal lines. The error bars represent one standard deviation extracted from 1000 bootstraps. **(G)** Dwell time distributions of SARS-CoV-2 polymerase activity traces at 25°C (gray circles) and 37°C (black circles) extracted from (B), and their respective fit to the pause-stochastic model (corresponding solid lines). **(H)** Nucleotide addition rate (green), Pause 1 (dark blue) and Pause 2 (cyan) exit rates at either 25°C or 37°C (solid and hatched bars, respectively) extracted from (G). The error bars in (C) and (G) represent one standard deviation extracted from 1000 bootstraps. The error bars in (H) are one standard deviation extracted from 100 bootstraps.

### 3’-dATP versus Remdesivir-TP: both ATP competitors but two different modes of incorporation

Next, we investigated how the elongation kinetics and the product length of SARS-CoV-2 polymerase were affected by two adenosine analogs, 3’-dATP and RDV-TP (**Fig. S2**). 3’-dATP is an obligatory terminator of RNA chain elongation for viral RdRp (*27*). RDV-TP has also been suggested to cause chain termination but only several cycles of nucleotide addition after its incorporation. If this is the true mechanism of action, then the experimental outcome of the presence of any of these two analogs should be indistinguishable in our assay.

In the presence of 500 µM NTP and 500 µM 3’-dATP, the ability of the SARS-CoV-2 polymerase to reach the end of the template (1043 nt) was compromised (**Fig. 2A** versus **Fig. 1B**). Indeed, increasing 3’-dATP concentration up to 2000 µM, only reduced the mean product length of SARS-CoV-2 polymerase by ∼1.7-fold, from (940 ± 13) *nt* to (566 ± 33) *nt* (mean ± standard deviation) (**Fig. 2B**), while not affecting the kinetics of RNA synthesis (**Fig. 2C, Fig. S3ABC**).

**Fig. 2.**
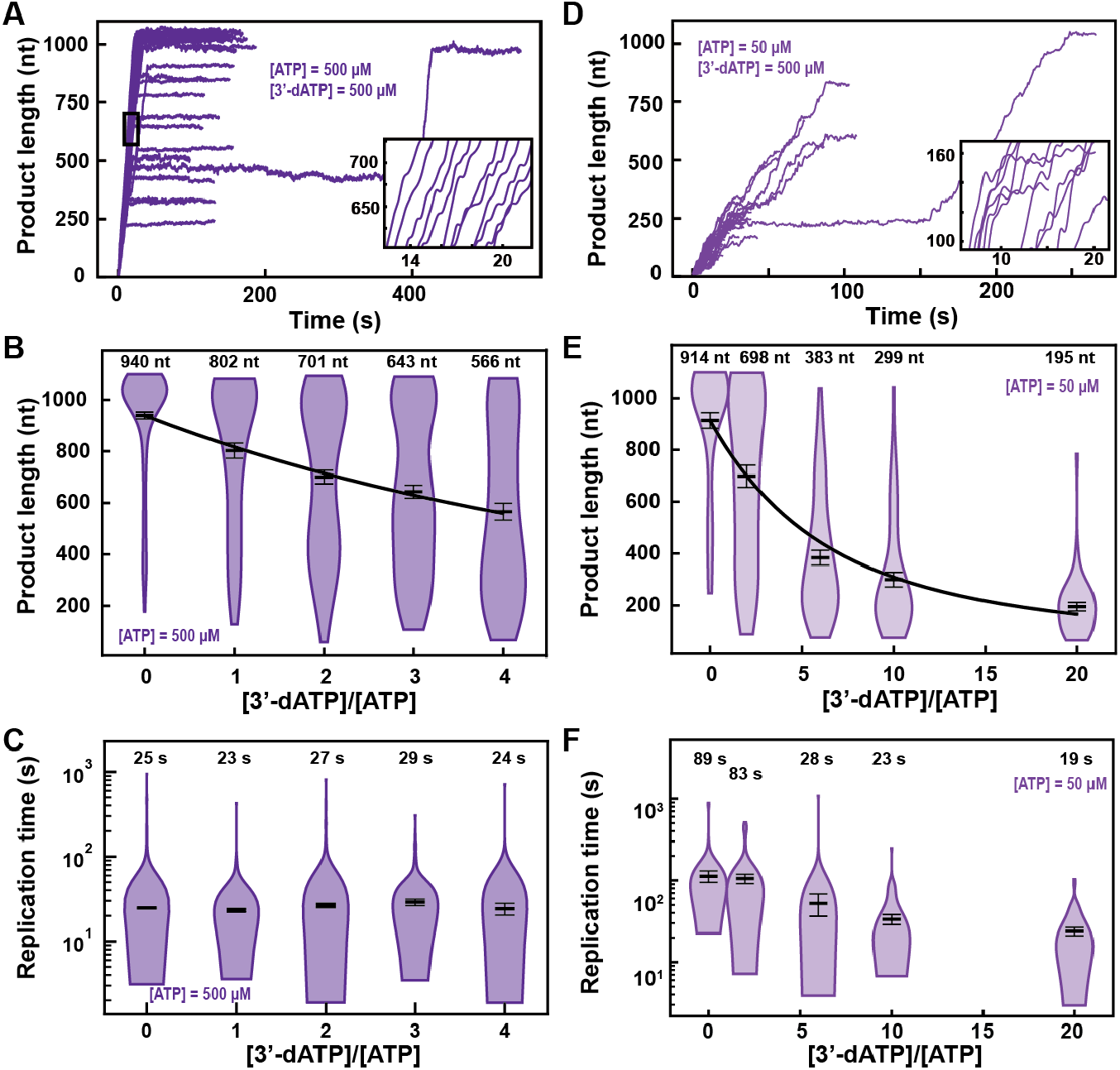
3’-dATP is an effective chain terminator for the SARS-CoV-2 polymerase. **(A)** SARS-CoV-2 replication traces for 500 µM NTPs and 500 µM 3’-dATP; and **(D)**, for 50 µM ATP, 500 µM all other NTPs and 500 µM 3’-dATP. **(B, E)** SARS-CoV-2 polymerase product length for the 1043 nt long template using the indicated concentration of ATP, 500 µM of other NTPs, as a function of [3’-dATP]/[ATP]. The mean values are indicated above the violin plots, and represented by horizontal black thick lines flanked by one standard deviation error bars extracted from 1000 bootstraps. **(C, F)** Replication time for the reaction conditions described in (B, E). The medians are indicated above the violin plots, and represented by horizontal black thick lines flanked by one standard deviation error bars extracted from 1000 bootstraps. In (B, E), the solid lines are the fits of the terminator effective incorporation rate (**Supplementary Materials**). In (A, D), the insets are zoom-in of the replication traces captured in the black square.

We derived a model to determine the effective incorporation rate *γ*, i.e. the average number of nucleotide addition cycles before terminator incorporation at equimolar concentration of competing natural nucleotide (in the presence of all NTPs) (**Supplementary Materials**). This model fits very well to the mean product length as a function of 3’-dATP:ATP stoichiometry (**Fig. 2B**), for example *γ*_3′*dATP*,50− µ*M ATP*_ = (780 ± 64) *nt*, meaning that the polymerase incorporates on average 780 nt before incorporating one 3’-dATP and terminating RNA synthesis (**Supplementary Materials**). A subsaturating concentration of NTP increases the probability to enter both the SNA and the VSNA pathways (**Fig. 1D**) (*25*), i.e. Pause 1 and Pause 2 probability, and would increase the effective incorporation rate of 3’-dATP, providing it is incorporated via any of the two slow nucleotide addition states. By decreasing ATP concentration from 500 to 50 µM, we indeed observed an increase in Pause 1 and Pause 2 probabilities by more than 2-fold and 3-fold, from (0.060 ± 0.002) to (0.149 ± 0.005) and from (0.0033 ± 0.0009) to (0.0115 ± 0.0026), respectively (**Fig. S3A-F**). Adding 500 µM of 3’-dATP significantly shortened the traces in comparison to the 500 µM ATP condition (**Fig. 2AD**). However, the effective incorporation rate of 3’-dATP was identical at both concentrations of ATP, i.e. *γ*_3′*dATP*,500− µ*M ATP*_ = (777 ± 50) *nt* (**Fig. 2E**), which indicates that 3’-dATP incorporation is only driven by stoichiometry, despite the significant increase in the SNA (Pause 1) and VSNA (Pause 2) pathways probabilities. Therefore, we conclude that 3’-dATP utilizes the NAB pathway for incorporation (**Fig. 1D**). Of note, the decrease in the median replication time is due to the shortening of the product length from early termination (**Fig. 2F**). Replicating the experiment at a 3’-dATP:ATP stoichiometry of 6 but now at 25 pN showed no significant differences in final product length in comparison to the data acquired at 35 pN (**Fig. 2E, Fig. S4A**).

RDV-TP is an adenine analog with a 1’-cyano modification that has recently been shown to outcompete ATP for incorporation (*17, 21*) (**Fig. S2**), while exhibiting a low cytotoxicity (*28*). RDV-TP has been proposed to induce delayed chain termination at i+3 (i being RDV incorporation position) during RNA synthesis by the core polymerase (*16, 17*). Adding 100 µM RDV-TP in a reaction buffer containing 500 µM NTPs showed a dramatic increase in the pause density and duration, but most of the traces reached the end of the template (**Fig. 3A**). We indeed observed a final product length largely unaffected at all concentrations of RDV-TP (**Fig. 3B**), while the median time for RNA synthesis increased by more than 10-fold (**Fig. 3C**), for RDV-TP concentrations increasing up to 300 µM. Therefore, the RDV-TP mechanism of action is not termination. We then investigated the origin of the pause induced by RDV-TP incorporation using our stochastic-pausing model (**Fig. 3D, Fig. S5B**). While the nucleotide addition rate is unaffected by RDV-TP, all pauses are significantly impacted. The exit rates of Pause 1 and Pause 2 decreased by 4- and 10-fold (**Fig. 3E**), while their probabilities increased by 2- and 4-fold, respectively (**Fig. 3F**). Most notably, the backtrack pause probability increased by 28-fold, from (0.0005 ± 0.0001) to (0.0142 ± 0.0015), when increasing RDV-TP concentration up to 300 µM. The backtrack pause probability increase was such that it most likely affected the probability and the exit rates of Pause 1 and Pause 2 above 50 µM RDV-TP (**Fig. 3F**).

**Fig. 3.**
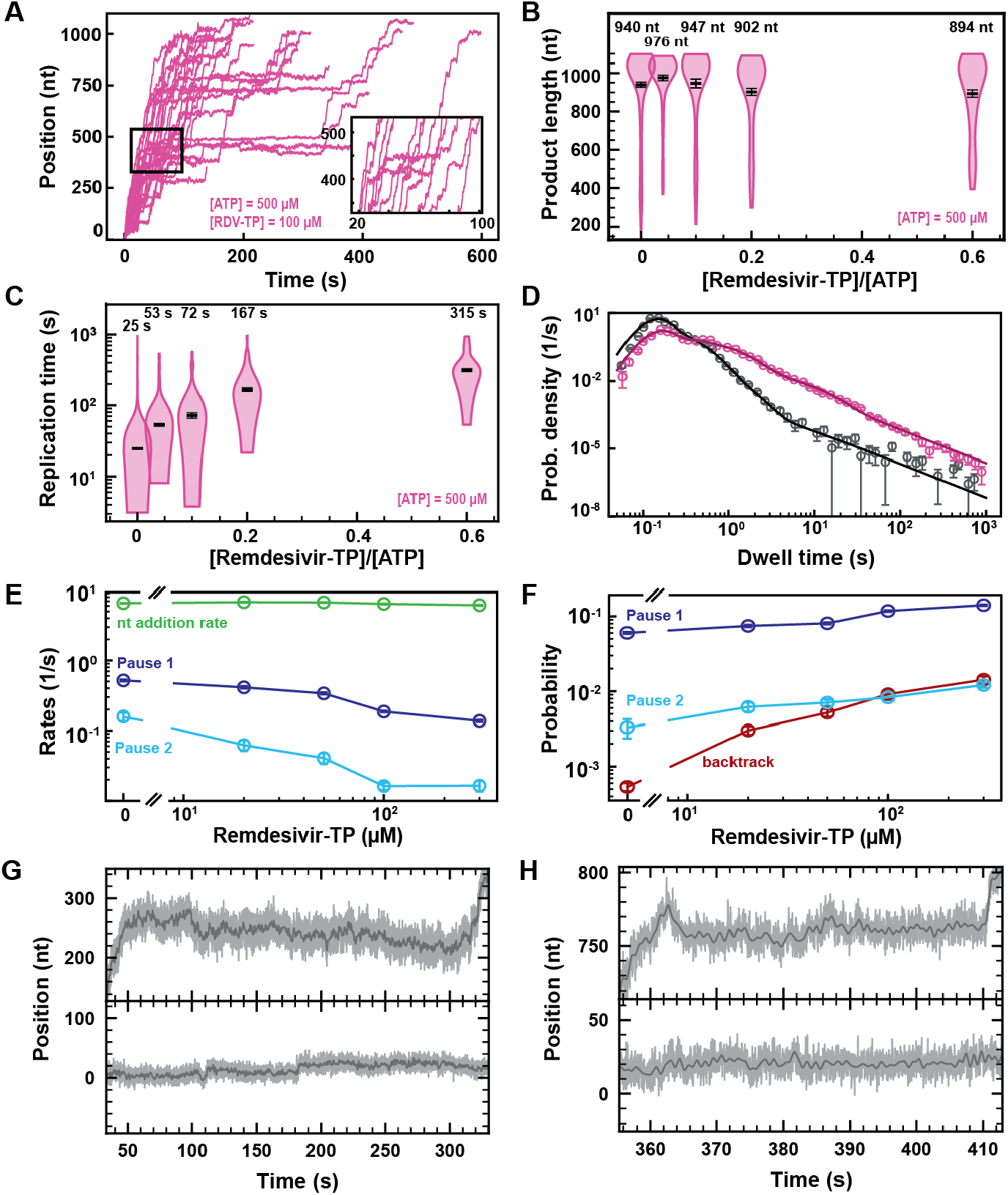
Remdesivir-TP (RDV-TP) is not a chain terminator but induces long-lived SARS-CoV-2 polymerase backtrack. **(A)** SARS-CoV-2 polymerase activity traces for 500 µM NTPs and 100 µM RDV-TP. The inset is a zoom-in of the polymerase activity traces captured in the black square. **(B)** SARS-CoV-2 polymerase product length for the 1043 nt long template using the indicated concentration of ATP, 500 µM of other NTPs, as a function of [RDV-TP]/[ATP]. The mean values are indicated above the violin plots, and represented by horizontal black thick lines flanked by one standard deviation error bars extracted from 1000 bootstraps. **(C)** Replication time for the reaction conditions described in (B). The median values are indicated above the violin plots, and represented by horizontal black thick lines flanked by one standard deviation error bars extracted from 1000 bootstraps. **(D)** Dwell time distributions of SARS-CoV-2 polymerase activity traces for 500 µM NTPs in the absence (gray) or presence of 100 µM RDV-TP (pink). The corresponding solid lines are the fit of the pause-stochastic model. **(E)** Nucleotide addition rate (green), Pause 1 (dark blue) and Pause 2 (cyan) exit rates for [NTPs] = 500 µM and several RDV-TP concentrations. **(F)** Probabilities to enter Pause 1 (dark blue), Pause 2 (cyan) and the backtrack (red) states for the conditions described in (E). Error bars in (D) represent one standard deviation extracted from 1000 bootstraps. Error bars in (E, F) are one standard deviation extracted from 100 bootstrap procedures. **(G, H)** Examples of deep SARS-CoV-2 backtracks induced by RDV-TP incorporation (top) and traces showing no polymerase activity (bottom). Traces acquired using an ultra-stable magnetic tweezers as described in Ref. (*25*) at 35 pN, 58 Hz acquisition frequency (grey), low-pass filtered at 1 Hz (dark grey), and using a SARS-CoV-2 polymerase reaction buffer containing 10 µM RDV-TP, 50 µM ATP and 500 µM all other NTPs.

As expected, the almost identical SARS-CoV-1 polymerase (*1*) demonstrated a similar kinetic signature to RDV-TP incorporation (**Fig. S6A-F, Table S1**), though to a lesser extent, e.g. the backtrack probability increased by ∼9-fold when raising RDV-TP concentration up to 300 µM versus 28-fold for SARS-CoV-2 polymerase.

To verify whether the applied tension modifies the incorporation kinetics of RDV-TP by the SARS-CoV-2 polymerase, we replicated the experiment using 500 µM NTP and 100 µM RDV-TP at 25 pN, i.e. a 10 pN lower force (**Fig. S4B**). We did not observe any significant difference between the two experiments at 35 and 25 pN (**Fig. S4CD**), indicating that tension does not play a significant role in RDV-TP incorporation.

Using our recently developed ultra-stable magnetic tweezers (*25*), we wanted to directly monitor polymerase backtrack induced by RDV-TP incorporation. To this end, we performed an experiment with 10 µM RDV-TP and 50 µM ATP, keeping all other NTPs at 500 µM (**Fig. 3GH, Fig. S5A**). We hypothesized that lowering the concentration of ATP, the natural competitor of RDV, would increase the incorporation yield of RDV and therefore polymerase backtrack probability. A close observation of the longest-lived pauses clearly demonstrate polymerase backtrack, as deep as ∼30 nt (**Fig. 3GH, Fig. S5A**), demonstrating that RDV incorporation induces polymerase backtrack, which leads to long-lived pauses.

To verify whether the incorporation of RDV-TP is stoichiometric, we further analyzed the experiment performed at 50 µM ATP, 500 µM all other NTPs, and 10 µM RDV-TP, at 25°C and 35 pN (**Fig. S7A**) (coined low ATP and RDV-TP concentrations), i.e. the same stoichiometry as 500 µM all NTPs and 100 µM RDV-TP (coined high ATP and RDV-TP concentrations). In absence of RDV-TP, the decrease in ATP concentration from 500 µM to 50 µM increased dramatically Pause 1 and Pause 2 probability by 2.5- and 23-fold, respectively, while the backtrack pause remained unchanged. The large increase in both Pause 1 and Pause 2 probabilities further disentangled the distribution of these pauses from the backtrack pause, and we therefore did not expect a strong crossover of the latter on the former (as observed at 500 µM all NTPs). We noticed an average product length of (633 ± 30) *nt* at low ATP and RDV-TP concentrations, i.e. a ∼30% shorter than for any other conditions presented in **Fig. S7B**, even though we acquired data for a much longer duration than at high ATP and RDV-TP concentrations, i.e. 11000 s vs. 1600 s, respectively. Interestingly, this result resembles what was observed at a 3’-dATP:ATP stoichiometry of ∼3 (**Fig. 2B**), indicating that RDV-TP induces what resembles termination at low ATP concentration. We also observed a ∼2.3-fold longer median replication time than at high ATP and RDV-TP concentrations (**Fig. S7C**), an increase largely underestimated as a large fraction of the traces never reached the end of the template during the measurement (**Fig. S7B**). Applying the stochastic-pausing model to the dwell time distribution of the low ATP and RDV-TP concentrations data (**Fig. S7D**), we found the nucleotide addition rate unchanged, while Pause 1 and Pause 2 exit rates were lower than in absence of RDV-TP, i.e. by 1.4- and 2.3-fold, respectively (**Fig. S7E**). At low ATP concentration, the probabilities of Pause 1 and Pause 2 were largely unaffected by the presence of RDV-TP, similarly to what was observed at 37°C (**Fig. S7F**). Most remarkably, the backtrack pause probability increased dramatically at low ATP and RDV-TP concentrations, even more so than at high ATP and RDV-TP concentrations, i.e. 43-fold versus 18-fold, respectively (**Fig. S7F**). The main effect of RDV-TP is to increase the backtrack pause probability. In our previous study of the impact of T-1106-TP on poliovirus RdRp, we showed that T-1106 incorporation induces long-lived backtrack pauses that appears as termination in ensemble assays (*22*). Interestingly, lowering ATP concentration increases the potency of RDV-TP by dramatically increasing the backtrack pause probability. However, we know these pauses are catalytically incompetent (*25*), and therefore the increase of the backtrack probability is an illustration of the effect of RDV-TP incorporation at low nucleotide concentration: the increased energy barrier induced by the steric clash of RDV-TP with the nsp12 serine-861 reduces dramatically the likelihood of a successful forward translocation of the polymerase (*17, 29*). This likelihood is even further reduced at low NTP concentration, which dramatically increases the probability of polymerase backtrack (*30*).

We previously observed that increasing the temperature helped to further disentangle the distributions of the different pauses (*24*). We therefore performed an experiment at 37°C in the presence of 100 µM RDV-TP and 500 µM all NTPs (**Fig. S8A**). The nucleotide addition rate significantly increased with temperature, while this increase was not affected by the presence of RDV-TP (**Fig. 1H, Fig. S8B**). On the one hand, Pause 1 and Pause 2 exit rates significantly decreased by 3- and 9-fold, respectively, when the reaction was performed with RDV-TP (**Fig. S8B**). On the other hand, Pause 1 and Pause 2 probabilities were unaffected by the presence of RDV-TP (**Fig. S8C**), supporting the notion that the increase in probability in the experiments performed at 25°C was the consequence of the polymerase backtrack pause distribution biasing Pause 1 and Pause 2 distributions (*22*). The backtrack pause probability still increased by more than 7-fold, i.e. from (0.0003 ± 0.0001) to (0.0022 ± 0.0007). The lesser increase in the backtrack pause probability at 37°C (28-fold at 25°C) is consistent with a model where RDV-MP represents a barrier to translocation, which crossing would be facilitated by increasing the thermal energy.

If RDV-TP incorporation resulted in a pause of similar exit rates as Pause 1 and Pause 2, but not mechanistically related to them, we would expect an increase in the probabilities of both pauses. However, in conditions where Pause 1 and Pause 2 distribution were clearly distinguishable from the backtrack pause distribution when having RDV-TP in the reaction buffer, i.e. at 37°C and at low ATP concentration, we did not observe an increase in Pause 1 and Pause 2 probabilities. Therefore, we suggest that RDV-TP is incorporated by the SNA and VSNA pathways (*25*), leading to polymerase backtrack when failing at overcoming the increased energy barrier resulting from the clash of RDV-MP with serine-861.

### T-1106-TP is incorporated with a low probability via the VSNA state

Pyrazine-carboxamides represent a promising family of antiviral NAs, of which the best-known member is Favipiravir (T-705), recently approved to treat influenza virus infection (*31*), and considered against SARS-CoV-2. We studied here another member of this family, T-1106 triphosphate (T-1106-TP), which is chemically more stable than T-705, while presenting similar antiviral properties (*22, 32*). T-1106-TP competes for incorporation against ATP and GTP in a sequence-dependent manner (*22, 32*). Adding 500 µM of T-1106-TP in a reaction buffer containing 500 µM NTPs significantly increased the number and duration of pauses observed in SARS-CoV-2 RNA synthesis activity traces (**Fig. 4A**), leading to a 2.6-fold increase in median replication time (**Fig. 4B**). For comparison, 50 µM of RDV-TP induced a median replication time as 500 µM T-1106-TP, at the same concentration of competing NTP, suggesting that RDV-TP is better incorporated than T-1106-TP. The final product length was not affected by T-1106-TP, consistent with the T-1106-TP mechanism of action not being termination (**Fig. 4C**) (*22*). Performing an experiment using the ultra-stable magnetic tweezers assay (*25*) with 500 µM T-1106 and 500 µM all NTPs at 35 pN force, the SARS-CoV-2 polymerase activity traces showed pauses with either a shallow backtrack (**Fig. S9A**), i.e. ≤10 nt, or no significant backtrack at all (**Fig. S9B**).

**Fig. 4.**
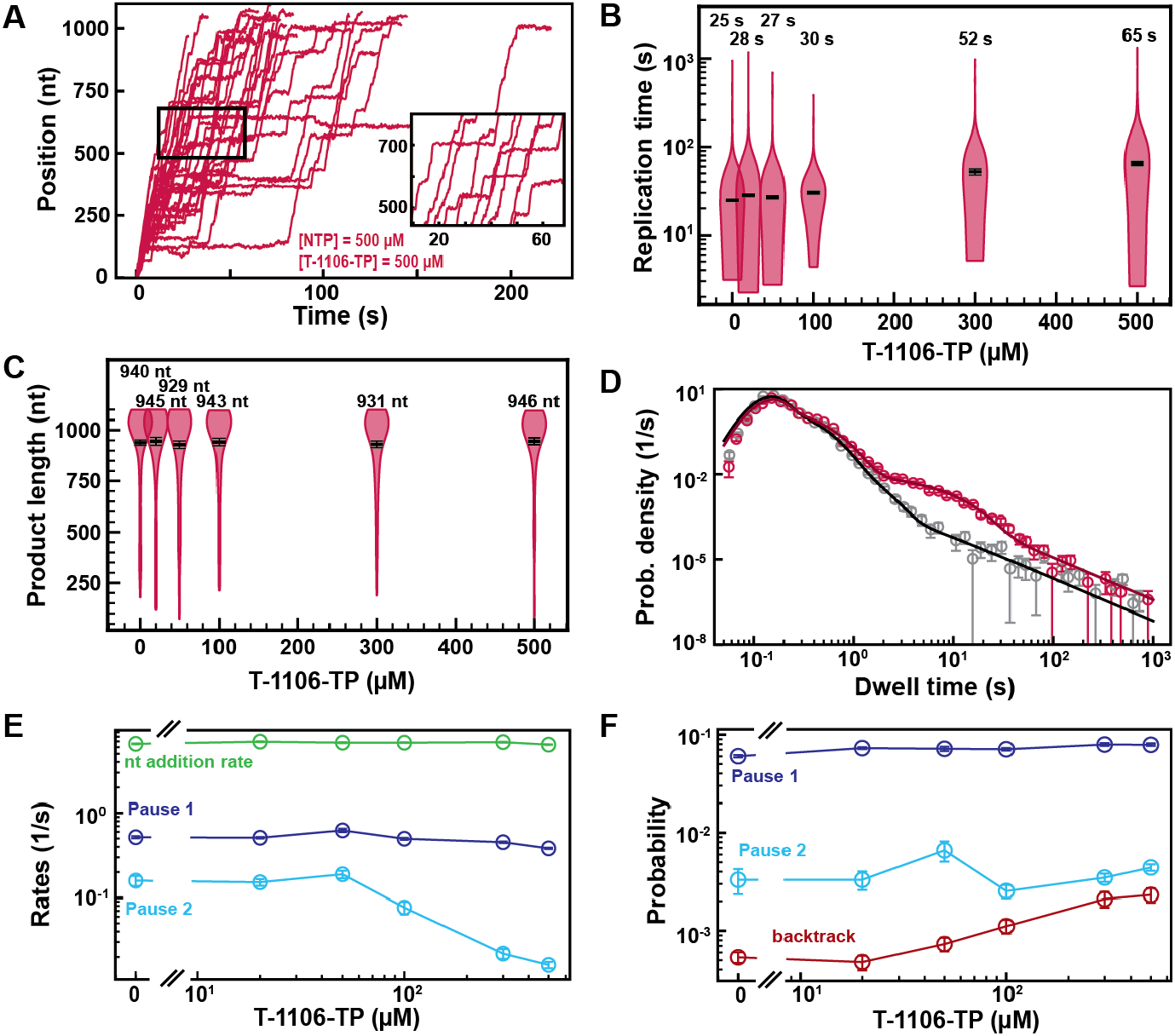
T-1106-TP incorporation induces pauses of intermediate duration and backtrack. **(A)** SARS-CoV-2 polymerase activity traces in the presence of 500 µM NTPs, in the presence of 500 µM T-1106-TP. The inset is a zoom-in of the polymerase activity traces captured in the black square. **(B)** SARS-CoV-2 replication time for the 1043 nt long template using 500 µM of all NTPs, and the indicated concentration of T-1106-TP. The median values are indicated above the violin plots, and represented by horizontal black thick lines flanked by one standard deviation error bars extracted from 1000 bootstraps. **(C)** SARS-CoV-2 polymerase product length using 500 µM NTPs and the indicated concentration of T-1106-TP. The mean values are indicated above the violin plots, and represented by horizontal black thick lines flanked by one standard deviation error bars extracted from 1000 bootstraps. **(D)** Dwell time distributions of SARS-CoV-2 polymerase activity traces for 500 µM NTP either without (gray) or with 500 µM (red) T-1106-TP. The corresponding solid lines are the fit to the pause-stochastic model. **(E)** Nucleotide addition rate (green), Pause 1 (dark blue) and Pause 2 (cyan) exit rates for [NTPs] = 500 µM and several T-1106-TP concentrations. **(F)** Probabilities to enter Pause 1 (dark blue), Pause 2 (cyan) and the backtrack (red) states for the conditions described in (E). The error bars denote one standard deviation from 1000 bootstraps in (D). Error bars in (E, F) are one standard deviation extracted from 100 bootstrap procedures.

Investigating how increasing T-1106-TP concentration affects SARS-CoV-2 RNA synthesis kinetics (**Fig. 4D, Fig. S9C**), we found that only the Pause 2 exit rate was affected, decreasing by 10-fold (**Fig. 4E**). Pause 1 and Pause 2 probabilities remained constant, while the backtrack pauses increased by almost 5-fold, though remaining in the low probability range, i.e. ∼0.002 (**Fig. 4F**). Repeating the experiment at 500 µM NTPs and 300 µM T-1106-TP at 25 pN, we found no difference in comparison to the data acquired at 35 pN (**Fig. S4E-G**). Here again, the tension has no significant effect. Our results suggest an incorporation of T-1106-TP only via the VSNA pathway (**Fig. 1D**), which explains its reduced promiscuity relative to RDV-TP, and is less likely than RDV-TP to induce polymerase backtrack upon incorporation. These observations contrast with our previous findings with poliovirus RdRp (*22*), where T-1106 incorporation induced deep polymerase backtrack. Therefore, the same NA may have a different mechanism of action on different RdRps.

### Sofosbuvir-TP is poorly incorporated by the SARS-CoV-2 polymerase

Next, we compared two uridine analogue chain terminators, i.e. Sofosbuvir and 3’-dUTP. Sofosbuvir presents a fluoro group at the 2’ α-position and a methyl group at the 2’ β-position, and is a non-obligatory chain terminator. Despite its low incorporation rate (*33*), Sofosbuvir has a proven antiviral effect against hepatitis C virus (HCV) and is an FDA-approved drug to treat HCV infection (*34, 35*). It is incorporated by SARS-CoV-2 polymerase (*17, 20*), but has no efficacy in infected cells (*36*). 3’-dUTP lacks a hydroxyl group in 3’ position, and is therefore an obligatory chain terminator.

The presence of 500 µM Sofosbuvir-TP with 500 µM NTP did not affect RNA synthesis by the SARS-CoV-2 polymerase (**Fig. 5A**), while early termination events appeared in the presence of 500 µM 3’-dUTP (**Fig. 5B**). Supporting this visual observation, the mean RNA product length of the SARS-CoV-2 polymerase was unaffected by the presence of Sofosbuvir-TP (**Fig. 5C**). Raising the 3’-dUTP:UTP stoichiometry to 4 reduced the mean product length by almost 5-fold, resulting in an effective incorporation rate *γ*_3′−*dUTP*,500 µ*M UTP*_ = (151 ± 6) *nt* (**Fig. 5D**). For both NAs, the replication time was unaffected (**Fig. S10A** and **S11A**), as well as SARS-CoV-2 RNA synthesis kinetics (**Fig. S10B-D** and **S11B-D**). Reducing the concentration of UTP down to 50 µM while keeping the other NTPs at 500 µM, Sofosbuvir-TP caused few early termination events when increased to 500 µM (**Fig. 5E**). Replacing Sofosbuvir-TP by 3’-dUTP, we observed a much stronger effect, as no activity traces reached the end of the template at 3’-dUTP:UTP stoichiometry of 10 (**Fig. 5F**). The analysis showed a limited impact of Sofosbuvir-TP on the mean product length, with a minimum of (563 ± 32) *nt* at a stoichiometry of 20 (**Fig. 5G**). 3’-dUTP was much more effectively incorporated, shortening the mean product length down to (67 ± 3) *nt* at the same stoichiometry (**Fig. 5H**). Their respective effective incorporation rate at 50 µM UTP reflected these observations, i.e. *γ*_*sofosbuvir*,50 µ*M UTP*_ = (3908 ± 467) *nt* and *γ*_3′−*dUTP*,50 µ*M UTP*_ = (241 ± 9) *nt*. In other words, SARS-CoV-2 polymerase incorporates on average 3908 nt and 241 nt before incorporating either a single Sofosbuvir-TP or a single 3’-dUTP, respectively. The kinetics of RNA synthesis were unaffected by the presence of either 3’-dUTP or Sofosbuvir-TP, while their median replication time decreased at high stoichiometry, a direct consequence of the shortening of the RNA synthesis product (**Fig. S10E-H** and **Fig. S11E-H**, respectively). Repeating the experiments for a Sofosbuvir-TP:UTP stoichiometry of 6 now at 25 pN tension (**Fig. S4A**), we did not see a significant difference in comparison with the data at 35 pN (**Fig. S5G**), therefore the applied tension has no influence in the incorporation of Sofosbuvir-TP. As for 3’-dATP, our data suggest that stoichiometry against the competing NTP regulates Sofosbuvir-TP and 3’-dUTP incorporation, which therefore support that these analogs utilize the NAB state pathway for incorporation (**Fig. 1D**). Our data provide further support to the poor incorporation of Sofosbuvir by SARS-CoV-2 (*17, 36*) and the low selectivity of the SARS-CoV-2 polymerase against 3’-dUTP.

**Fig. 5.**
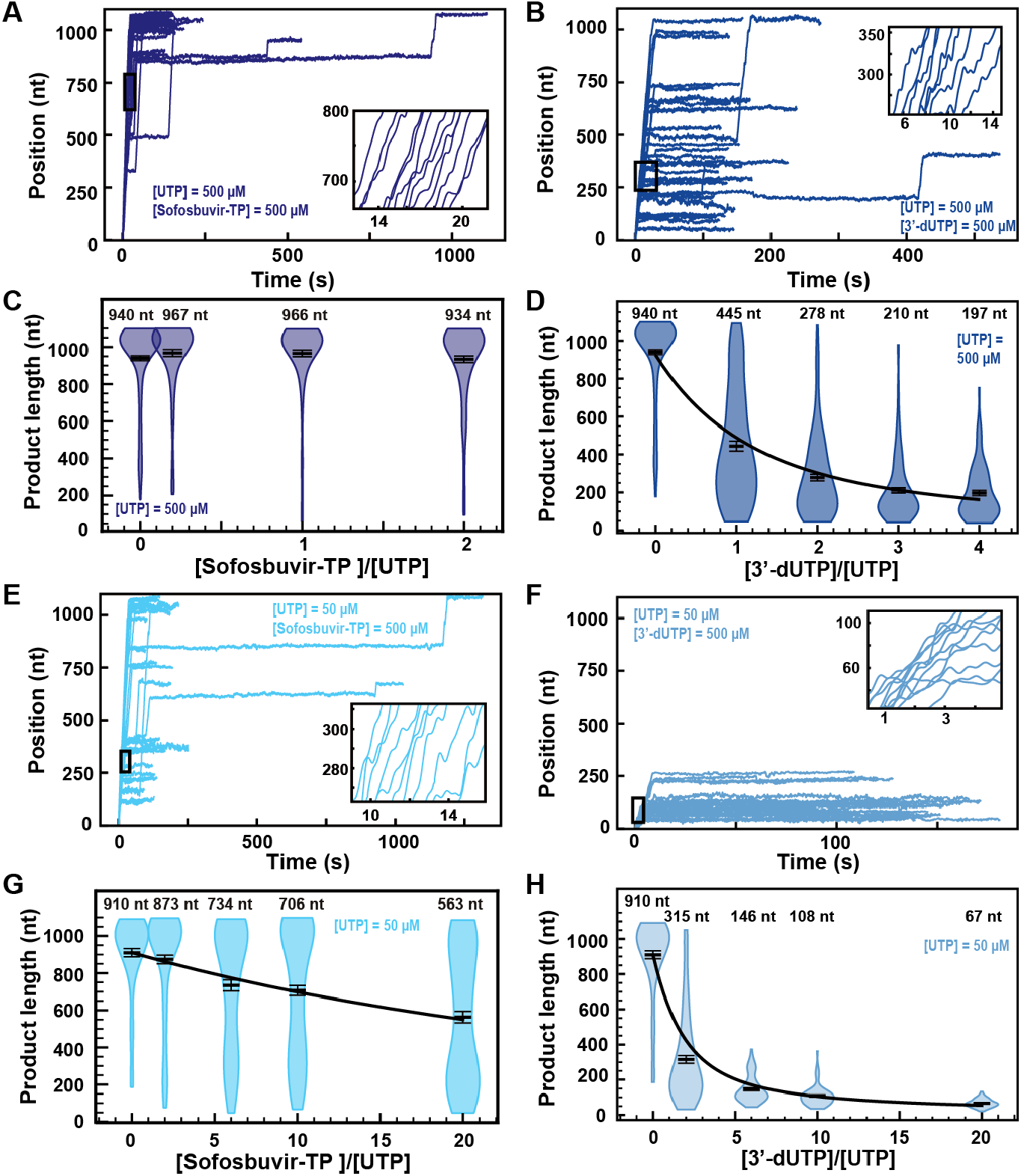
Sofosbuvir-TP is a poor SARS-CoV-2 polymerase inhibitor in contrast with 3’-dUTP. **(A, B)** SARS-CoV-2 polymerase activity traces for 500 µM NTPs and 500 µM of either (A) Sofosbuvir-TP or (B) 3’-dUTP. **(C, D)** SARS-CoV-2 polymerase product length using the indicated concentration of UTP, 500 µM of other NTPs, as a function of either (C) [Sofosbuvir-TP]/[UTP] or (D) [3’-dUTP]/[UTP]. The mean values are indicated above the violin plots, and represented by horizontal black thick lines flanked by one standard deviation error bars extracted from 1000 bootstraps. **(E, F)** SARS-CoV-2 polymerase activity traces in the presence 50 µM of UTP, 500 µM of all other NTPs and 500 µM of either (E) Sofosbuvir-TP or (F) 3’-dUTP. **(G, H)** SARS-CoV-2 polymerase product length using 50 µM UTP, 500 µM of other NTPs, as a function of either (G) [Sofosbuvir-TP]/[UTP] or (H) [3’-dUTP]/[UTP]. The mean values are indicated above the violin plots, and represented by horizontal black thick lines flanked by one standard deviation error bars extracted from 1000 bootstraps. In (D, G, H), the solid line is the fit of the terminator effective incorporation rate (**Supplementary Materials**). In (A, B, E, F), the insets are a zoom-in of the replication traces captured in the black square.

### ddhCTP is well incorporated by the polymerase but does not affect SARS-CoV-2 replication in cells

3′-Deoxy-3′,4′-didehydro-CTP (ddhCTP) is a recently discovered natural antiviral NA produced in mammalian cells by the viperin-catalyzed conversion of CTP to ddhCTP using a radical-based mechanism (*37*). While ddhCTP has been shown to efficiently terminate flavivirus replication both *in vitro* and in cells, its antiviral activity against SARS-CoV-2 polymerase remains unknown. The addition of 500 µM ddhCTP to a reaction buffer containing 500 µM NTP induces early termination events in the SARS-CoV-2 polymerase activity traces (**Fig. 6A**). Similar amount of 3’-dCTP instead of ddhCTP resulted in a larger fraction of traces showing early termination events (**Fig. 6B**). The average RNA product length of the SARS-CoV-2 polymerase decreased by 1.4-fold when raising the ddhCTP:CTP stoichiometry to 4 (**Fig. 6C**), while it decreased by 2.7-fold at similar stoichiometry against CTP (**Fig. 6D**). We measured a respective effective incorporation rate *γ*_*ddhCTP*,500 µ*M CTP*_ = (1221 ± 130) *nt* and *γ* _3′−*dUTP*,500 µ*M CTP*_ = (338 ± 18) *nt* (**Fig. 6CD**). For both NAs, the replication time (**Fig. S12A** and **S13A)** and the RNA synthesis kinetics (**Fig. S12B-D** and **S13B-D**) were largely unaffected. Reducing the concentration of CTP down to 50 µM and keeping the other NTPs at 500 µM, both ddhCTP and 3’-dCTP showed a significant reduction in length of the activity traces (**Fig. 6EF**). Analyzing the average product length, we extracted the respective effective incorporation rates at 50 µM CTP, i.e. *γ*_*ddhCTP*,50 µ*M CTP*_ = (1360 ± 71) *nt* and *γ*_3′−*dCTP*,50 µ*M CTP*_ = (457 ± 21) *nt* (**Fig. 6GH)**. These values are similar as what was measured at 500 µM CTP, and confirms the better incorporation of 3’-dCTP over ddhCTP. The kinetics of RNA synthesis were unaffected by the presence of ddhCTP or 3’-dCTP, while their median replication time decreased at high stoichiometry, as a result of the shortening of the RNA synthesis product (**Fig. S12F-H** and **S13F-H**, respectively). We also did not observe any impact of the applied tension for ddhCTP incorporation (**Fig. S4A**). Here again, stoichiometry against their competing natural nucleotide CTP directly dictates the incorporation of 3’-dCTP and ddhCTP, further supporting the utilization of the NAB pathway for their incorporation (**Fig. 1D**).

**Fig. 6.**
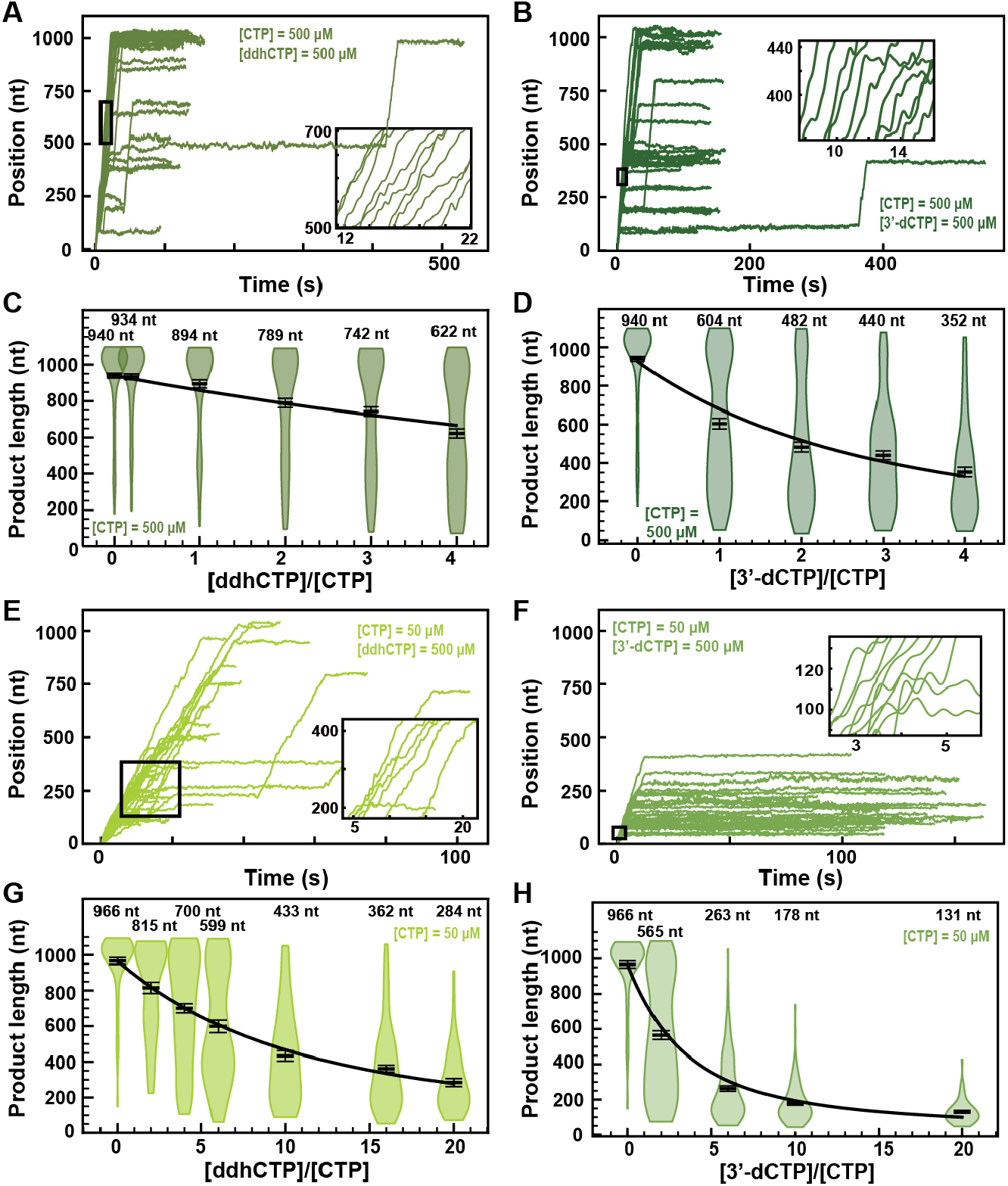
ddhCTP and 3’-dCTP inhibit efficiently the SARS-CoV-2 polymerase. **(A, B)** SARS-CoV-2 polymerase activity traces for 500 µM NTPs and 500 µM of either (A) ddhCTP or (B) 3’-dCTP. **(C, D)** SARS-CoV-2 polymerase product length using the indicated concentration of CTP, 500 µM of other NTPs, as a function of either (C) [ddhCTP]/[CTP] or (D) [3’-dCTP]/[CTP]. The mean values are indicated above the violin plots, and represented by horizontal black thick lines flanked by one standard deviation error bars extracted from 1000 bootstraps. **(E, F)** SARS-CoV-2 polymerase activity traces in the presence 50 µM of CTP, 500 µM of all other NTPs and 500 µM of either (E) ddhCTP or (F) 3’-dCTP. **(G, H)** SARS-CoV-2 polymerase activity traces product length using 50 µM CTP, 500 µM of other NTPs, as a function of the stoichiometry of either (G) [ddhCTP]/[CTP] or (H) [3’-dCTP]/[CTP]. The mean values are indicated above the violin plots, and represented by horizontal black thick lines flanked by one standard deviation error bars extracted from 1000 bootstraps. In (C, D, G, H), the solid lines are the fits of the terminator effective incorporation rate (**Supplementary Materials**). In (A, B, E, F), the insets are a zoom-in of the replication traces captured in the black square.

Though not as high as 3’-dCTP, the effective incorporation rate of ddhCTP should be sufficient to demonstrate a certain efficacy against viral replication in cells. Indeed, ddhCTP is a chain terminator, therefore a single incorporation is sufficient to end RNA synthesis. To verify whether ddhCTP inhibits replication in cells, we infected Huh7-hACE2 cells with SARS-CoV-2, treated these cells with different concentrations of RDV, Sofosbuvir and ddhC, and report on the level of infection by immunofluorescence against SARS-CoV-2 N protein (**Supplementary Materials, Fig. S14AB**). While RDV showed a clear antiviral effect with an EC_50_ of 0.007 µM (**Fig. S14B**), ddhC and Sofosbuvir did not show any impact on SARS-CoV-2 replication in cells. This result suggests that SARS-CoV-2 is able to evade the antiviral properties of the endogenously synthesized antiviral nucleotide analog ddhC. We hypothesized that the 3’ to 5’ exonuclease activity of nsp14 protects SARS-CoV-2 replication by excising ddhCMP from the nascent RNA. To test this hypothesis, we made a SARS-CoV-2 strain, which includes the amino acid substitutions D90A and E92A that remove the exoribonuclease activity of nsp14 (**Fig. S15A**). This SARS-CoV-2 mutant was unable to replicate in cells (**Fig. S15B-F**), confirming a recent report (*38*). Therefore, the role of nsp14 in removal of ddhCMP from the SARS-CoV-2 genome could not be verified experimentally. Future experiments will be designed to address this question.

## Discussion

We present here the first characterization of the mechanism of action of antiviral nucleotide analogs against SARS-CoV-2 polymerase at the single-molecule level. We show that SARS-CoV-2 polymerase is the fastest RNA studied polymerase to date, elongating up to ∼170 *nt*. *s*^−1^ at 37 °C (**Fig. 1H**). With our assay, we monitored the incorporation and determined the mechanism of action of several NAs, i.e. 3’-dATP, 3’-dUTP, 3’-dCTP, Sofosbuvir-TP, ddhCTP, T-1106-TP and RDV-TP.

Our study definitely proves that RDV-TP is not a chain terminator at physiological concentration of all NTPs, but instead induces pauses in the polymerase elongation kinetics that are easily overcome at saturating NTP concentration (**Fig. 3**). This finding has been corroborated by two recent studies since our preprint was published on BioRxiv in August 2020 (*29, 39*). Similarly, T-1106-TP incorporation does not induce termination, but pauses in the polymerase elongation kinetics (**Fig. 4**). However, RDV-TP affects both Pause 1 and Pause 2 exit rates, while T-1106-TP affects only the latter. We showed here that these two NAs do not affect the probability to enter Pause 1 and Pause 2, suggesting that they preferably bind to the polymerase active site after it entered the SNA (Pause 1) or VSNA (Pause 2) pathway. Indeed, if the pauses induced by either RDV-TP or T-1106-TP incorporation were mechanistically unrelated to Pause 1 and Pause 2, the total number of pauses would cumulate and the probability of pausing would dramatically increase, which we do not observe. We therefore suggest that RDV-TP can be incorporated by both SNA and VSNA pathways, while T-1106-TP is only incorporated by the latter. Finally, Pause 1 and Pause 2 respectively account for ∼6% and ∼0.3% of all the nucleotide addition events at a saturating concentration of NTP. This defines an upper limit for RDV-TP and T-1106-TP relative incorporation, and explains why RDV-TP is incorporated much better than Favipiravir (*40*).

Two recent ensemble kinetic studies investigating the mechanism of action of RDV-TP on SARS-CoV-2 elongation kinetics have recently been published. In the first one, the experiments were performed at submicromolar concentration of NTPs, and showed that RDV-TP is incorporated 3-fold better than ATP in such condition (*17*). In the second one, the authors also claimed that RDV-TP was better incorporated than ATP (*21*), while using higher concentration of NTPs than in the first study. However, in both studies, the substrate concentration was subsaturating, a condition in which the SNA and VSNA probabilities significantly increase (*25*), favoring RDV-TP incorporation. Importantly, none of these studies monitored the incorporation of RDV-TP in competition with saturating concentration of ATP, and are therefore not representative of the in vivo conditions (*41*). Last, the authors of the second study extracted the RDV-TP incorporation kinetics through several successive rounds of RDV-TP incorporation (*21*). Not only is this scenario highly unlikely to happen in the cell, it may also bias the measured kinetics. Our work does not present any of these biases, as we were able to monitor RDV-TP incorporation in competition with saturating concentration of NTP – including ATP –, while the SARS-CoV-2 polymerase was elongating a ∼1 kb long RNA product.

Our assay revealed that RDV-TP incorporation leads the coronavirus polymerase into backtrack as deep as ∼30 nt (**Fig. 3GH**). This result demonstrates that the barrier induced by the clash of RDV-MP (*29*) with the serine-861 of nsp12 is sufficiently strong to elicit polymerase backtrack, leading the polymerase into a pause long enough to be mistaken for a termination event in ensemble assays. We anticipate that RDV efficacy is further amplified when the polymerase is elongating through template secondary structures, which stimulates polymerase backtrack (*25*). Lower ATP concentration would also decrease the probability to overcome the barrier when an uracil is encoded ∼3 nt downstream the incorporated RDV-MP, increasing the backtrack pause probability, as observed here. Interestingly, RDV has a strong efficacy against SARS-CoV-2 in infected cells (**Fig. S14AB**), which indicates that the 3’ to 5’ exonuclease nsp14 does not remove efficiently RDV-MP from the nucleic acid chain. Our results suggest that polymerase backtrack is therefore not an intermediate of product strand proofreading, which corroborates a preceding study showing that nsp14 cannot excise single stranded RNA (*9*).

Concerning obligatory terminators, the effective incorporation rate we measured showed that 3’-dATP (**Fig. 2**), 3’-dUTP (**Fig. 5**), 3’-dCTP (**Fig. 6**), and – to a lesser extent – ddhCTP (**Fig. 6**) are well incorporated by the SARS-CoV-2 polymerase, while Sofosbuvir-TP is strongly outcompeted by UTP (**Fig. 5**). Though well incorporated, 3’-dNTP are cytotoxic, and are therefore not used as antiviral drugs (*42*). Interestingly, the effective incorporation rate of all these terminators is only affected by the stoichiometry of their respective competing natural nucleotide, and not their absolute concentration (unlike RDV-TP), suggesting an incorporation via the NAB pathway (**Fig. 1D**). Indeed, we showed that NA incorporated via either the SNA or the VSNA pathway, e.g. RDV-TP, would be more likely to be added in the RNA chain at low substrate concentration, independently of the stoichiometry.

A steady-state kinetic study showed that nucleotide analogs modified at the 2’ and 3’ positions are strongly discriminated against by their competing natural nucleotide (*17*). Such selectivity is an issue for purine-based analogs, which must compete with high concentrations of ATP and GTP in the cell. In contrast, pyrimidine-based analogs, for example derivatives of CTP, will only need to compete with intracellular CTP pools on the order of 100 µM (*41*). These features make ddhCTP a particularly attractive antiviral NA. Furthermore, under certain conditions, the interferon α-induced viperin converts up to 30% of the cellular pool of CTP into ddhCTP, further increasing the ddhCTP:CTP stoichiometry in a direction favoring even greater potency (*37*). However, we could not show any efficacy of ddhC in SARS-CoV-2 infected cells (**Fig. S14**), suggesting that SARS-CoV-2 has developed ways to counter this cellular defense mechanism. Future studies will investigate whether the exonuclease nsp14 is capable of removing ddhCMP and is therefore responsible for protecting the virus against endogenously produced antiviral NAs.

High-throughput, real-time magnetic tweezers present numerous advantages to study RdRp elongation dynamics over a discontinuous assay, and therefore demand integration of such an assay into any pipeline investigating the selectivity and/or mechanism of action of NAs. Such an assay will also reveal how adding functional capacity to the core polymerase, for example RNA helicase activity and proofreading, modulate RdRp elongation dynamics and response to antiviral therapeutics.

## Materials and Methods

### High-throughput magnetic tweezers apparatus

The high-throughput magnetic tweezers used in this study have been described in detail elsewhere (*43*). Shortly, a pair of vertically aligned permanent magnets (5 mm cubes, SuperMagnete, Switzerland) separated by a 1 mm gap are positioned above a flow cell (see paragraph below) that is mounted on a custom-built inverted microscope. The vertical position and rotation of the magnets are controlled by two linear motors, M-126-PD1 and C-150 (Physik Instrumente PI, GmbH & Co. KG, Karlsruhe, Germany), respectively. The field of view is illuminated through the magnets gap by a collimated LED-light source, and is imaged onto a large chip CMOS camera (Dalsa Falcon2 FA-80-12M1H, Stemmer Imaging, Germany) using a 50× oil immersion objective (CFI Plan Achro 50 XH, NA 0.9, Nikon, Germany) and an achromatic doublet tube lens of 200 mm focal length and 50 mm diameter (Qioptic, Germany). To control the temperature, we used a system described in details in Ref. (*24*). Shortly, a flexible resistive foil heater with an integrated 10MΩ thermistor (HT10K, Thorlabs) is wrapped around the microscope objective and further insulated by several layers of kapton tape (KAP22-075, Thorlabs). The heating foil is connected to a PID temperature controller (TC200 PID controller, Thorlabs) to adjust the temperature within ∼0.1°.

### Flow cell assembly

The fabrication procedure for flow cells has been described in details in Ref. (*43*). To summarize, we sandwiched a double layer of Parafilm by two #1 coverslips, the top one having one hole at each end serving as inlet and outlet, the bottom one being coated with a 0.01% m/V nitrocellulose in amyl acetate solution. The flow cell is mounted into a custom-built holder and rinsed with ∼1 ml of 1x phosphate buffered saline (PBS). 3 µm diameter polystyrene reference beads are attached to the bottom coverslip surface by incubating 100 µl of a 1:1000 dilution in PBS of (LB30, Sigma Aldrich, stock conc.: 1.828*10^11^ particles per milliliter) for ∼3 minutes. The tethering of the magnetic beads by the RNA hairpin construct relies on a digoxygenin/anti-digoxygenin and biotin-streptavidin attachment at the coverslip surface and the magnetic bead, respectively. Therefore, following a thorough rinsing of the flow cell with PBS, 50 µl of anti-digoxigenin (50 µg/ml in PBS) is incubated for 30 minutes. The flow cell was flushed with 1 ml of 10 mM Tris, 1 mM EDTA pH 8.0, 750 mM NaCl, 2 mM sodium azide buffer to remove excess of anti-digoxigenin followed by rinsing with another 0.5 ml of 1x TE buffer (10 mM Tris, 1 mM EDTA pH 8.0 supplemented with 150 mM NaCl, 2 mM sodium azide). The surface is then passivated by incubating bovine serum albumin (BSA, New England Biolabs, 10 mg/ml in PBS and 50% glycerol) for 30 minutes, and rinsed with 1x TE buffer.

### Single molecule RdRp replication activity experiments

20 µl of streptavidin coated Dynal Dynabeads M-270 streptavidin coated magnetic beads (ThermoFisher) was mixed with ∼0.1 ng of RNA hairpin (total volume 40 µl) (**Supplementary Materials**) and incubated for ∼5 minutes before rinsing with ∼2 ml of 1x TE buffer to remove any unbound RNA and the magnetic beads in excess. RNA tethers were sorted for functional hairpins by looking for the characteristic jump in extension length due to the sudden opening of the hairpin during a force ramp experiment (**Fig. S1B**) (*44*). The flow cell was subsequently rinsed with 0.5 ml reaction buffer (50 mM HEPES pH 7.9, 2 mM DTT, 2 µM EDTA, 5 mM MgCl_2_). After starting the data acquisition at a force that would keep the hairpin open, 100 µl of reaction buffer containing either 0.6 µM of nsp12, 1.8 µM of nsp7 and nsp8 for SARS-CoV-2 experiments or 0.1 µM of nsp12, 1 µM of nsp7 and nsp8 for SARS-CoV-1 experiments, the indicated concentration of NTPs and of nucleotide analogs (if required) were flushed in the flow cell to start the reaction. Sofosbuvir-TP and T-1106-TP were purchased from Jena Bioscience (Jena, Germany) and 3’-dATP was purchased from TriLink Biotechnologies (San Diego, CA, USA). The experiments were conducted at a constant force as indicated for a duration of 20 to 40 minutes. The camera frame rate was fixed at either 58 Hz or 200 Hz, for reaction temperature set at either 25°C or 37°C, respectively. A custom written Labview routine (*45*) controlled the data acquisition and the (x-, y-, z-) positions analysis/tracking of both the magnetic and reference beads in real-time. Mechanical drift correction was performed by subtracting the reference bead position to the magnetic bead position.

### Data processing

The replication activity of SARS-CoV-2 core polymerase converts the tether from ssRNA to dsRNA, which concomitantly decreases the end-to-end extension of the tether. The change in extension measured in micron was subsequently converted into replicated nucleotides *N_R_* using the following equation:

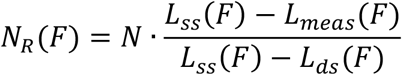

Where, *L_meas_*(*F*), *L_ss_*(*F*) and *L_ds_*(*F*) are the measured extension during the experiment, the extension of a ssRNA and of a dsRNA construct, respectively, experiencing a force *F*, and *N* the number of nucleotides of the ssRNA template (*23*). The traces were then filtered using a Kaiser-Bessel low-pass filter with a cut-off frequency at 2 Hz. As previously described in Ref. (*23*), a dwell time analysis was performed by scanning the filtered traces with non-overlapping windows of 10 nt to measure the time (coined throughout the manuscript dwell time) for SARS-CoV-2 polymerase to incorporate ten successive nucleotides. The dwell times of all the traces for a given experimental condition were assembled and further analyzed using a maximum likelihood estimation (MLE) fitting routine to extract the parameters from the stochastic-pausing model (**Supplementary Materials**).

### SARS-CoV-2 replication product length analysis

To extract the product length of the replication complex, only the traces where the beginning and the end could clearly be distinguished and for which the tether did not rupture for ten minutes following the last observed replication activity were considered. We represented the mean product length, as well as one standard deviation of the mean from 1000 bootstraps as error bars.

## Author contribution

DD and CEC conceived the research. MS, SCB and DD performed and analyzed the single molecule experiments, PvN wrote the MLE analysis routine. FSP made the RNA construct used for the study. MD derived the terminator competition model. LDH, JMW, SCA and TG provided ddhCTP. BC, TTNL and AS provided SARS-CoV-1 proteins. RNK provided SARS-CoV-2 proteins. XM and YX performed the in cells SARS-CoV-2 replication assay in the presence of nucleotide analogs. HX and PYS performed the in cells SARS-CoV-2 exonuclease mutant replication experiment. DD and CEC wrote the article. All the authors have discussed the data and edited the article.

## Acknowledgements

We thank Joy Feng from Gilead Sciences for providing RDV-TP, and Veronique Fattorini and Barbara Selisko for excellent technical assistance and help in SARS-CoV-1 proteins purification. DD was supported by the Interdisciplinary Center for Clinical Research (IZKF) at the University Hospital of the University of Erlangen-Nuremberg, the German Research Foundation grant DFG-DU-1872/3-1 and BaSyC – Building a Synthetic Cell” Gravitation grant (024.003.019) of the Netherlands Ministry of Education, Culture and Science (OCW) and the Netherlands Organisation for Scientific Research (NWO). DD thanks OICE for providing office and lab space, and access to their molecular biology lab. RNK was supported by grant AI123498 from NIAID, NIH. JMW and LDH thank the Ministry of Business Innovation & Employment Contract UOOX1904 (NZ). Y.X was supported by the National Institutes of Health (NIH) grant AI151638. SARS-Related Coronavirus 2, Isolate USA-WA1/2020 (NR-52281) was deposited by the Centers for Disease Control and Prevention and obtained through BEI Resources, NIAID, NIH. PYS was supported by NIH grants AI134907 and UL1TR001439, and awards from the Sealy & Smith Foundation, Kleberg Foundation, the John S. Dunn Foundation, the Amon G. Carter Foundation, the Gilson Longenbaugh Foundation, and the Summerfield Robert Foundation. JJA and CEC were supported by grant AI045818 from NIAID, NIH. AS, TTNL and BC acknowledge grants by the Fondation pour la Recherche Médicale (Aide aux équipes), the SCORE project H2020 SC1-PHE-Coronavirus-2020 (grant#101003627), and the REACTing initiative (REsearch and ACTion targeting emerging infectious diseases).

