## Supplementary Materials for "Inhibition of SARS-CoV-2 polymerase by nucleotide analogs: a single molecule perspective"

**This PDF file includes:**

Supplementary text

Fig. S1 to S15

Table S1 and S2

**Supplementary text**

*ddhCTP production*

ddhCTP was prepared as previously described (manuscript in preparation). Briefly, ddhC (*1*) was dissolved in 20 mM Tris-HCl, 100 mM KCl, 10 mM BME at pH 7.5. ATP was added to a final concentration of 100 µM, and PEP was added to a concentration of ~3 mM. The proteins human UCK2, CMPK1, and NDK were all added to the reaction mixture to a final concentration of ~10 µM. PK/LDH mixture were added at a final concentration of 1.2 and1.8 units•$mL^{-1}$. After the reaction was complete, proteins were precipitated by lowering the pH to 2 with concentrated HCl and then immediately returning the pH to 9. Precipitated protein was removed by centrifugation and the supernatant was passed through a 0.22 µm filter. The final solution was diluted 10-fold using 20 mM TEAB at pH 9.5. ddhCTP was purified with a MonoQ 5/50 anion exchange column using TEAB buffer at pH 9.5. The final ddhCTP was concentrated with lyophilization. Concentration of ddhCTP stocks were determined using an extinction coefficient of 9,000 $M^{-1}$•$cm^{-1}$.

*Recombinant Protein Expression of RdRp (nsp12) and cofactors (nsp7 and nsp8) from SARS-CoV-2*

This protocol was described in Ref. (*2*). SARS-CoV-2 nsp12: The SARS-CoV-2 nsp12 gene was codon optimized and cloned into *pFastBac* with C-terminal additions of a TEV site and strep tag (Genscript). The pFastBac plasmid and DH10Bac E. coli (Life Technologies) were used to create recombinant bacmids. The bacmid was transfected into Sf9 cells (Expression Systems) with Cellfectin II (Life Technologies) to generate recombinant baculovirus. The baculovirus was amplified through two passages in Sf9 cells, and then used to infect 1 L of Sf21 cells (Expression Systems) and incubated for 48 h at 27°C. Cells were harvested by centrifugation, resuspended in wash buffer (25 mM HEPES pH 7.4, 300 mM NaCl, 1 mM $MgCl_2$, 5 mM DTT) with 143 µl of BioLock per liter of culture. Cells were lysed via microfluidization (Microfluidics). Lysates were cleared by centrifugation and filtration. The protein was purified using Strep Tactin superflow agarose (IBA). Strep Tactin eluted protein was further purified by size exclusion chromatography using a Superdex 200 Increase 10/300 column (GE Life Sciences) in 25 mM HEPES, 300 mM NaCl, 100 µM $MgCl_2$, 2 mM TCEP, at pH 7.4. Pure protein was concentrated by ultrafiltration prior to flash freezing in liquid nitrogen. *SARS-CoV-2 nsp7 and nsp8:* The SARS-CoV-2 nsp7 and nsp8 genes were codon optimized and cloned into pET46 (Novagen) with an N-terminal 6x histidine tag, an enterokinase site, and a TEV protease site. Rosetta2 pLys E. coli cells (Novagen) were used for bacterial expression. After induction with isopropyl β-D-1-thiogalactopyranoside (IPTG), cultures were grown at 16°C for 16 h. Cells were harvested by centrifugation and pellets were resuspended in wash buffer (10 mM Tris pH 8.0, 300 mM NaCl, 30 mM imidazole, 2 mM DTT). Cells were lysed via microfluidization and lysates were cleared by centrifugation and filtration. Proteins were purified using Ni-NTA agarose beads and eluted with wash buffer containing 300 mM imidazole. Eluted proteins dialyzed into dialysis buffer (10mM Tris pH 8.0, 300mM NaCl, 2mM DTT) with 1% w/w TEV protease at room temperature overnight. Digested proteins were passed back over Ni-NTA agarose beads. Digested proteins were further purified by size exclusion chromatography using a Superdex 200 Increase 10/300 column (GE Life Sciences). Purified proteins were concentrated by ultrafiltration prior to flash freezing with liquid nitrogen.

*Recombinant Protein Expression of RdRp (nsp12) and cofactors (nsp7 and nsp8) from SARS-CoV-1*

This protocol was described in Ref. (*3*). All SARS-CoV proteins used in this study were expressed in Escherichia coli (E. coli), under the control of T5 promoters. Cofactors nsp7L8 and nsp8 alone were expressed from pQE30 vectors with C-terminal and N-terminal hexa-histidine tags respectively. TEV cleavage site sequences were included for His-tag removal following expression. The nsp7L8 fusion protein was generated by inserting a GSGSGS linker between nsp7- and nsp8-coding sequences. Cofactors were expressed in NEB Express C2523 (New England Biolabs) cells carrying the pRare2LacI (Novagen) plasmid in the presence of Ampicillin (100 µM/mL) and Chloramphenicol (17 µg/mL). Protein expression was induced with 100 µM IPTG once the OD600 = 0.5–0.6, and expressed overnight at 17°C. Protein was purified first through affinity chromatography with HisPur Cobalt resin (Thermo Scientific), with a lysis buffer containing 50 mM Tris-HCl pH 8, 300 mM NaCl, 10 mM Imidazole, supplemented with 20 mM $MgSO_4$, 0.25 mg/mL Lysozyme, 10 µg/mL DNase, 1 mM PMSF, with lysis buffer supplemented with 250 mM imidazole. Eluted protein was concentrated and dialyzed overnight in the presence of histidine labeled TEV protease (1:10 w/w ratio to TEV:protein) for removal of the protein tag. Cleaved protein was purified through a second cobalt column and protein was purified through size exclusion chromatography (GE, Superdex S200) in gel filtration buffer (25 mM HEPES pH 8, 150 mM NaCl, 5mM $MgCl_2$, 5 mM TCEP). Concentrated aliquots of protein were flash-frozen in liquid nitrogen and stored at -80°C. A synthetic, codon-optimized SARS-CoV nsp12 gene (DNA 2.0) bearing C-terminal 8His-tag preceded by a TEV protease cleavage site was expressed from a pJ404 vector (DNA 2.0) in E. coli strain BL21/pG-Tf2 (Takara). Cells were grown at 37°C in the presence of Ampicillin and Chloramphenicol until OD600 reached 2. Cultures were induced with 250 µM IPTG and protein expressed at 17°C overnight. Purification was performed as above in lysis buffer supplemented with 1% CHAPS. Two additional wash steps were performed prior to elution, with buffer supplemented with 20 mM imidazole and 50 mM arginine for the first and second washes respectively. Polymerase was eluted using lysis buffer with 500 mM imidazole and concentrated protein was purified through gel filtration chromatography (GE, Superdex S200) in the same buffer as for nsp7L8. Collected fractions were concentrated and supplemented with 50% glycerol final concentration and stored at -20 °C.

*Cell lines and viruses*

Huh7 cells expressing human ACE2 (huh7-hACE2) were established by transducing huh7 cells with lentiviral particles derived with pWPI-IRES-Puro-Ak-ACE2 (a gift from Sonja Best; Addgene plasmid # 154985). SARS-CoV-2, isolate USA-WA1/2020 (NR-52281), was obtained through BEI Resources and propagated once on VERO E6 cells before it was used for this study.

*Immunofluorescence assay*

Huh7-hACE2 cells in 96-well plates (Corning) were infected with SARS-CoV-2 (USA-WA1/2020 isolate) at MOI of 0.05 in DMEM supplemented with 1% FBS. 1.5 h before the viral inoculation, the tested compounds were added to the wells in triplicate. The infection proceeded for 24 h without the removal of the viruses or the compounds. The cells were then fixed with 4% paraformaldehyde, permeabilized with 0.1% Triton-100, blocked with DMEM containing 10% FBS, and stained with a rabbit monoclonal antibody against SARS-CoV-2 NP (GeneTex, GTX635679) and an Alexa Fluor 488-conjugated goat anti-mouse secondary antibody (ThermoFisher Scientific). Hoechst 33342 was added in the final step to counterstain the nuclei. Fluorescence images of approximately ten thousand cells were acquired per well with a 10x objective in a Cytation 5 (BioTek). The total number of cells, as indicated by the nuclei staining, and the fraction of the infected cells, as indicated by the NP staining, were quantified with the cellular analysis module of the Gen5 software (BioTek).

*SARS-CoV-2 virus production and characterization*

SARS-CoV-2 WT and nsp14 exoribonuclease knockout viruses were prepared using a SARS-CoV-2 infectious clone (*4*). Briefly, viral RNA was obtained by in vitro RNA transcription, and 40 µg RNA transcripts and 20 µg N gene RNA were co-electroporated into $8 \times 10^6$ Vero E6 cells using Gene Pulser XCell electroporation system (Bio-Rad, Hercules, CA) at a setting of 270 V and 950 µF with a single pulse. The electroporated cells were seeded to a T75 flask and immediately transfer to BSL-3 facility. Viral production was confirmed by RT-PCR. The supernatants of electroporated cells were harvested and centrifuged at 1,000 g for 10 min to remove cell debris. 250 µl supernatant was added and mixed thoroughly with 1 ml of TRIzol LS reagent (Thermo Fisher Scientific). RNA was extracted according to the manufacture's instruction and resuspended in 20 µl of nuclease-free water. RT-PCR was performed using the SuperScript® IV One-Step RT-PCR kit (Thermo Fisher Scientific).

Virus was determined by plaque assay. Approximately $1.2\times10^6$ Vero E6 cells were seeded to each well of a 6-well plate. The viruses were 10-fold serially diluted with 2% FBS DMEM medium and 200 µl of virus dilution was transferred to each well of the 6-well plate. After the incubation for 1 h at 37°C, 2 ml of overlay medium containing 2% FBS DMEM medium and 1% sea-plaque agarose (Lonza, Walkersville, MD), was added to the infected cells per well. After a 2-day incubation, another 2 ml of overlay medium with neutral red (final concentration 0.01%) was added onto the first overlay. After 12 h incubation, the plates were sealed with Breath-Easy sealing membrane (Sigma-Aldrich, St. Louis, MO) and plaques were counted.

*SARS-CoV-2 luciferase replicon assay*

SARS-CoV-2 transient luciferase replicon assay was performed as previously described (*5*). WT and mutant replicon RNA, and N gene mRNA were obtained through T7 *in vitro* transcription, and 40 µg RNA transcripts and 20 µg N gene RNA were co-electroporated into $8\times10^6$ Huh-7 cells using Gene Pulser XCell electroporation system (Bio-Rad) at a setting of 270V and 950 µF with a single pulse. After 10 min recovery, electroporated cells were seeded to 24-well plates, and harvested at indicated timepoints. Luciferase signal was measured using *Renilla* luciferase assay system (Promega) and read by Cytation 5 (Bio Tek) according to the manufacturer's protocols.

*Construct fabrication.*

The fabrication of the RNA hairpin has been described in detail in Ref. (*6*). The RNA hairpin is made of a 499 bp double-stranded RNA stem terminated by a 20 nt loop that is assembled from three ssRNA annealed together, and two handles, one of 856 bp at the 5' end and one 822 bp at the 3' end. The handles include either a 343 nt digoxygenin-labeled ssRNA or a 443 nt biotin-labeled ssRNA. Upon applied force above ~21 pN, the hairpin opens and frees a 1043 nt ssRNA template for SARS-CoV-2 replication. To obtain the different parts of the RNA construct, template DNA fragments were amplified via PCR, purified (Monarch PCR and DNA cleanup kit) and in vitro transcribed (NEB HiScribe™ T7 High Yield RNA Synthesis Kit). Transcripts were then treated with Antarctic Phosphatase and T4 Polynucleotide Kinase. RNAs were purified using the RNA Clean & Concentrator-25 kit (Zymo Research). Individual RNA fragments were annealed and ligated with T4 RNA ligase 2 (NEB) to assemble the RNA hairpin.

The template contains 250 U (24%), 253 A (24%), 273 C (26%) and 267 G (26%).

*Stochastic-pausing model*

The model is described in detail in (*7-9*). There are many kinetic models that are consistent with the empirical dwell-time distributions we observe, and we here work under the assumption that the probability of pausing is low enough that there is only one rate-limiting pause in each dwell-time window. This assumption washes out most details of the kinetic scheme that connects pauses and nucleotide addition, but allows us to determine the general form of the dwell-time distribution without specifying how the pauses are connected to the nucleotide addition pathway

$$p_{dw}(t) \propto p_{na}\Gamma\left(t; N_{dw}, \frac{1}{k_{na}}\right) + Q(t)\left(\sum_{n=1}^{N_{sp}} p_n k_n e^{-k_n t} + \frac{a_{bt}}{2(1+t/1s)^{3/2}}\right). \qquad \text{(Eq. 1)}$$

In the above expression, the gamma function in the first term contributes the portion $p_{na}$ of dwell times that originate in the RdRp crossing the dwell time window of size $N_{dw}$ base pairs without pausing; the second term is a sum of contributions originating in pause-dominated transitions, each contributing a fraction $p_n$ of dwell times; the third term captures the asymptotic power-law decay (amplitude $a_{\mathrm{bt}}$) of the probability of dwell-times dominated by a backtrack. The backtracked asymptotic term needs to be regularized for times shorter than the diffusive backtrack step. We have introduced a regularization at 1 s, but the precise timescale does not matter, as long as it is set within the region where the exponential pauses dominate over the backtrack. From left to right, each term of Equation 1 is dominating the distribution for successively longer dwell-times.

A cut-off factor $Q(t)$ for short times is introduced to account for the fact that the dwell time window includes $N_{dw}$ nucleotide-addition steps,

$$Q(t) = \frac{(tk_{na}/N_{dw})^{N_{dw}-1}}{1+(tk_{na}/N_{dw})^{N_{dw}-1}}.$$

The fit results dependence on these cut-offs is negligible as long as they are introduced in regions where the corresponding term is sub-dominant. Here the cut is placed under the center of the elongation peak, guaranteeing that it is placed where pausing is sub-dominant.

*Maximum likelihood estimation*

The normalized version of Equation 1 is the dwell time distribution fit to the experimentally collected dwell-times $\{t_i\}_i$ by minimizing the likelihood function (*10*)

$$L = -\sum_i \ln p_{dw}(t_i) \qquad \text{(Eq. 2)}$$

with respect to rates and probabilistic weights.

*Dominating in a dwell-time window vs. dominating in one step*

The fractions $p_n$ represent the probability that a particular rate $k_n$ dominates the dwell-time. We want to relate this to the probability $P_n$ that a specific exit rate dominates within a one-nt transcription window. Assuming we have labelled the pauses so that $k_{n-1} > k_n$, we can relate the probability of having rate $n$ dominating in $N_{dw}$ steps to the probability of having it dominate in one step through

$$p_n = \left(\sum_{m=0}^{n} P_m\right)^{N_{dw}} - \left(\sum_{m=0}^{n-1} P_m\right)^{N_{dw}}, \quad p_0 = p_{na} = P_{na}^{N_{dw}} = P_0^{N_{dw}} \qquad \text{(Eq. 3)}$$

The first term in Equation 3 represents the probability of having no pauses longer than the $n^{th}$ pause in the dwell-time window, and the second term represents the probability of having no pauses longer than the $(n-1)^{th}$ pause. The difference between the two terms is the probability that the $n^{th}$ pause will dominate. This can be inverted to yield a relation between the single-step probabilities ($P_n$) and the dwell-time window probabilities ($p_n$)

$$P_n = \left(\sum_{m=0}^{n} p_m\right)^{1/N_{dw}} - \left(\sum_{m=0}^{n-1} p_m\right)^{1/N_{dw}}, \qquad P_0 = p_0^{1/N_{dw}}$$

This relationship has been used throughout the manuscript to relate our fits over a dwell time window to the single-step probabilities.

*Maximum likelihood estimation (MLE) fitting routine*

The above stochastic-pausing model was fit to the dwell time distributions using a custom Python 3.7 routine. Shortly, we implemented a combination of simulated annealing and bound constrained minimization to find the parameters that minimize Equation 2. We calculated the statistical error on the parameters by applying the MLE fitting procedure on 100 bootstraps of the original data set (*11*), and reported the standard deviation for each fitting parameters.

*Competition between obligatory terminator nucleotide analogues and their natural nucleotide homologues*

Starting with an empty active site (E), we assume that there is direct binding competition between the natural nucleotide (N) and the nucleotide analogue terminator (T, simply coined terminator) that result in either the former bound (Nb) or the latter bound (Tb) to the active site. From these states there can be any number of intermediate states before the base is either added to the chain with probability $P_{\mathrm{cat}}^{\mathrm{T/N}}$, or unbinds from the pocket with probability $1-P_{\mathrm{cat}}^{\mathrm{T/N}}$ (see figure below).

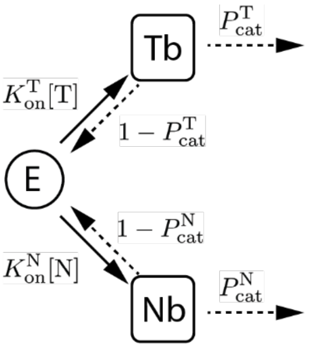


From the empty active site state (E), either a terminator (T) or a natural nucleotide (N) can bind though direct competition with the first order binding rates $K_{\mathrm{on}}^{\mathrm{T}}[\mathrm{T}]$ and $K_{\mathrm{on}}^{\mathrm{N}}[\mathrm{N}]$ (solid arrows represent rates) respectively. From the bound states (Tb/Nb) there can be many sub steps before either incorporating the base with probability $P_{\mathrm{cat}}^{\mathrm{T/N}}$, or ejecting it from the active site with probability and $1-P_{\mathrm{cat}}^{\mathrm{T/N}}$ (dashed arrows represent probabilities).

The effective incorporation rate is the attempt rate times the probability of success,

$$k_{\mathrm{inc}}^{\mathrm{T/N}} = [\mathrm{T/N}] K_{\mathrm{on}}^{\mathrm{T/N}} P_{\mathrm{cat}}^{\mathrm{T/N}}$$

and the relative probability that next incorporated base is a terminator or natural nucleotide is given by the relative effective addition rates

$$\frac{p^{\mathrm{T}}}{p^{\mathrm{N}}} = \frac{k_{\mathrm{inc}}^{\mathrm{T}}}{k_{\mathrm{inc}}^{\mathrm{N}}} = \frac{[\mathrm{T}]}{[\mathrm{N}]}\frac{K_{\mathrm{on}}^{\mathrm{T}}}{K_{\mathrm{on}}^{\mathrm{N}}}\frac{P_{\mathrm{cat}}^{\mathrm{T}}}{P_{\mathrm{cat}}^{\mathrm{N}}}, \qquad p^{\mathrm{T}} + p^{\mathrm{N}} = 1.$$

This can be rewritten as

$$p^{\mathrm{N}} = \frac{\gamma}{\gamma + x}, \qquad p^{\mathrm{T}} = \frac{x}{\gamma + x}, \qquad x = \frac{[\mathrm{T}]}{[\mathrm{N}]}, \qquad \gamma = \frac{K_{\mathrm{on}}^{\mathrm{N}}}{K_{\mathrm{on}}^{\mathrm{T}}}\frac{P_{\mathrm{cat}}^{\mathrm{N}}}{P_{\mathrm{cat}}^{\mathrm{T}}}$$

In the above $x$ is the relative stoichiometry between T and N, while $\gamma$ is the relative effective incorporation rates of N and T at equimolar conditions.

On an infinite construct, polymerization will proceed until the first T is incorporated, after which it terminates. At termination, the product has incorporated $n-1$ Ns, and finally one T, with probability

$$P(n) = \left(p^{\mathrm{N}}\right)^{n-1} p^{\mathrm{T}} = \left(1 - p^{\mathrm{T}}\right)^{n-1} p^{\mathrm{T}}.$$

The average number of Ns and Ts incorporated on an infinite construct is therefore

$$n^{\infty} = \sum_{n=1}^{\infty} n \left(p^{\mathrm{N}}\right)^{n-1} p^{\mathrm{T}} = 1/p^{\mathrm{T}}.$$

If the construct only allows for the addition of $N$ Ns and Ts, the average number of Ns and Ts in the product will instead be

$$n^N = \sum_{n=1}^{N} n_\mathrm{A} \left(p^\mathrm{N}\right)^{n-1} p^\mathrm{T} + \sum_{n=N+1}^{\infty} N \left(p^\mathrm{N}\right)^{n-1} p^\mathrm{T} = \frac{1-\left(p^\mathrm{N}\right)^N}{p^\mathrm{T}} = n^\infty \left(1-\left(p^\mathrm{N}\right)^N\right).$$

For a genome of length $L$, with the relative abundance $q$ of templating bases for N and T, we thus expect there to be at most $N = qL$ Ns and Ts incorporated at termination. At termination the product then has the average length

$$l^L = \frac{n^{qL}}{q} = \frac{1-\left(p^\mathrm{N}\right)^{qL}}{q\,p^\mathrm{T}} = l^\infty \left(1-\left(p^\mathrm{N}\right)^{qL}\right), \qquad l^\infty = \frac{1}{q\,p^\mathrm{T}}$$

*Data fitting*

Though the constructs are 1043 nucleotides long, this length is not always reached even when there are no terminators in the buffer. The average product length is about 10% shorter than the full construct length. To account for this reduction in maximal average product length, we simply fix $L$ to be the mean product length reached without terminator in the buffer, and fit out $\gamma$ from a least square fit, weighted with the inverse experimental variance.

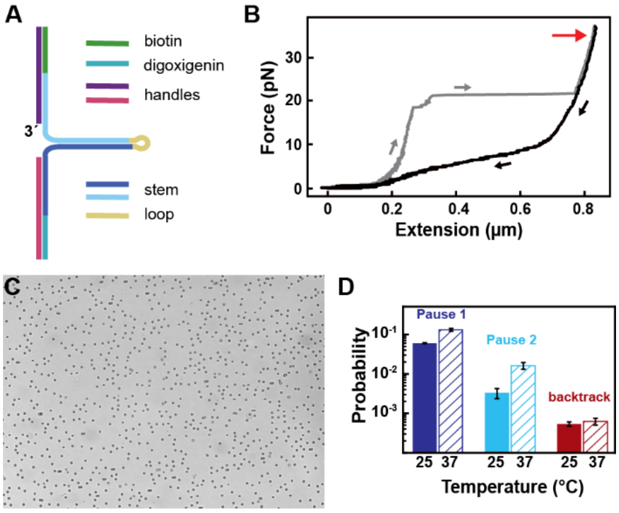


**Figure S1: Experimental conditions and data representation of SARS-CoV-2 high throughput magnetic tweezers experiments. (A)** Schematic of the RNA hairpin construct assembled from hybridizing and ligating single stranded RNAs. **(B)** Force as a function of the extension of the RNA hairpin presented in (A). Increasing and decreasing force ramp represented in gray and black, respectively. The red arrow indicates the extension at 35 pN, the force applied in the measurements unless specified. **(C)** Typical field of view containing ~450 hairpin tethered magnetic beads in a high throughput magnetic tweezers assay. **(D)** Probabilities to enter Pause 1 (dark blue), Pause 2 (cyan) and the backtrack (red) states at either 25°C (solid bars) or 37°C (hatched bars). Error bars in (D) are one standard deviation extracted from 100 bootstraps.

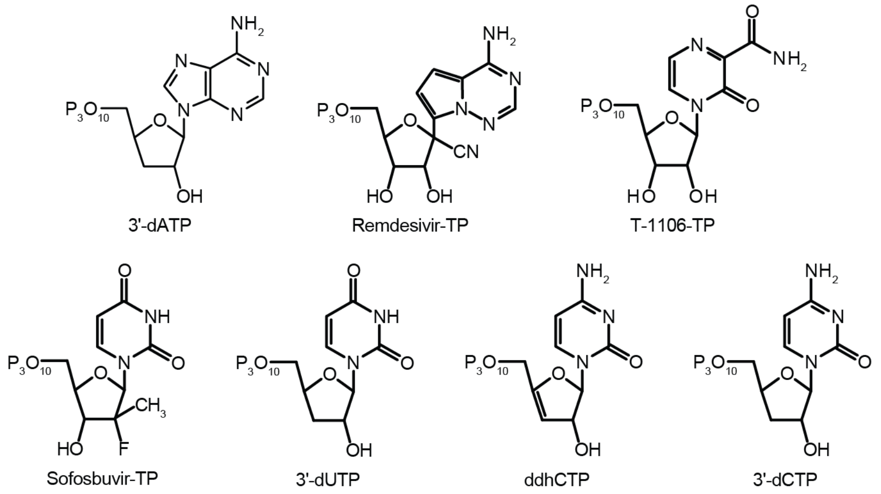


**Figure S2: Structure of the nucleotide analogues used in this study.**

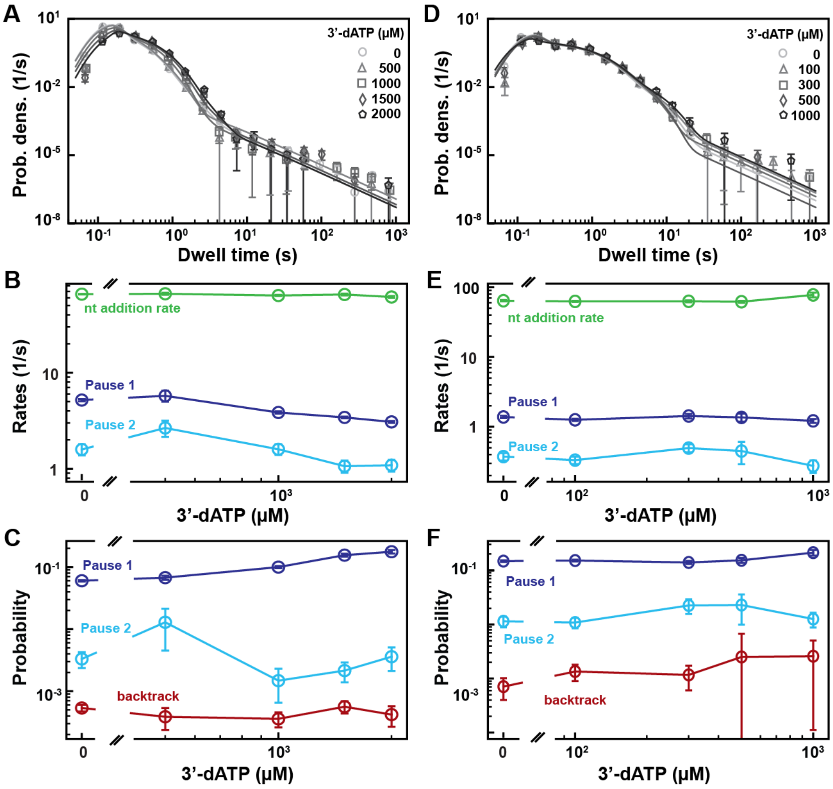


**Figure S3: SARS-CoV-2 polymerase activity traces kinetics in presence of 3'-dATP. (A)** Dwell time distributions of SARS-CoV-2 polymerase activity traces acquired in the presence of 500 µM NTP either without (circles) or with 0.5 mM (triangles), 1 mM (squares), 1.5 mM (diamonds), 2 mM (pentagons) 3'-dATP. The color code from light to dark gray further highlights the increasing concentration of 3'-dATP. The solid lines represent the fit to the pause-stochastic model. **(B)** Nucleotide addition rate (green), Pause 1 (dark blue) and Pause 2 (cyan) exit rates for the conditions described in (A). **(C)** Probabilities to enter Pause 1 (dark blue), Pause 2 (cyan) and the backtrack (red) states for the conditions described in (A). **(D)** Dwell time distributions of SARS-CoV-2 polymerase activity traces acquired in the presence of 50 µM ATP, 500 µM of other NTPs, and either without (circles), or with 0.1 mM (triangles), 0.3 mM (squares), 0.5 mM (diamonds) and 1 mM (pentagons) 3'-dATP. The color code from light to dark gray further highlights the increasing concentration of 3'-dATP. The solid lines represent the fit of the pause-stochastic model. **(E)** Nucleotide addition rate (green), Pause 1 (dark blue) and Pause 2 (cyan) exit rates for the conditions described in (D). **(F)** Probabilities to enter Pause 1 (dark blue), Pause 2 (cyan) and the backtrack (red) states for the conditions described in (D). **(**The error bars in (A and D) represent one standard deviation extracted from 1000 bootstraps. The error bars in (B, C, E, F) are one standard deviation extracted from 100 bootstrap.

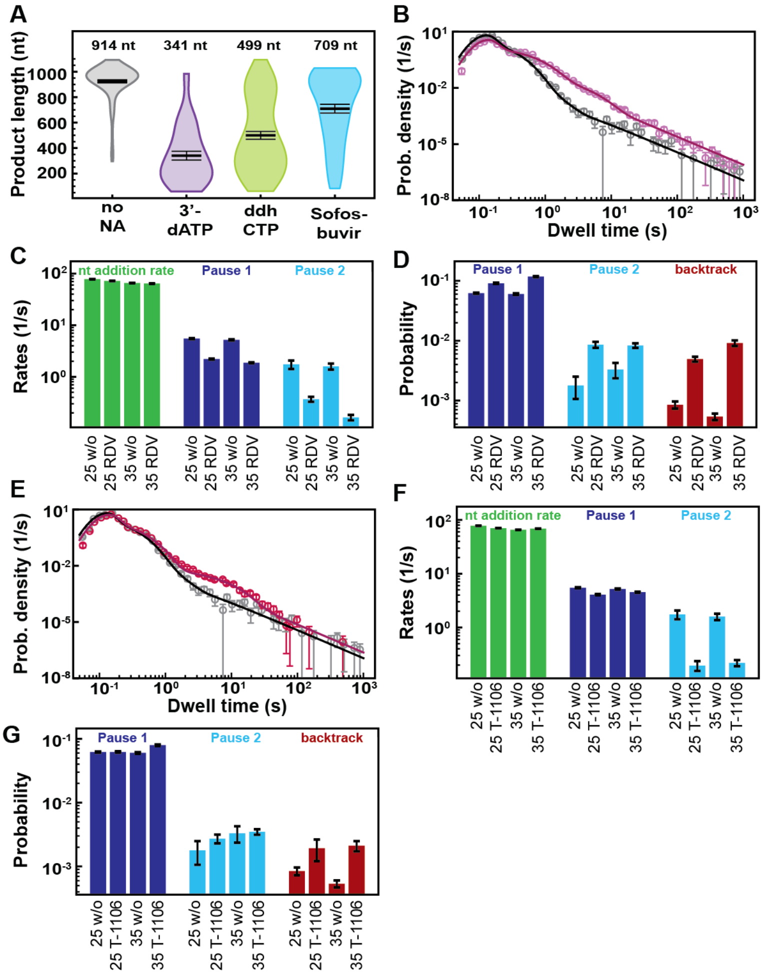


**Figure S4: Decreasing the applied tension does not change the effect of nucleotide analogs on the SARS-CoV-2 polymerase elongation.** All measurements are done at 25°C. **(A)** Product length of SARS-CoV-2 polymerase at 50 µM ATP and 500 µM all other NTPs (gray) or 300 µM nucleotide analog and 50 µM of the competing NTP (purple: 3'-dATP, green: ddhCTP, blue: Sofosbuvir-TP). The mean values are indicated above the violin plots, and represented by horizontal black thick lines flanked by one standard deviation error bars extracted from 1000 bootstraps. **(B)** Dwell time distributions of SARS-CoV-2 polymerase activity traces for 500 µM all NTPs without (gray) or with 100 µM RDV-TP (pink) at 25 pN. The corresponding solid lines are the fit to the pause-stochastic model. **(C)** Nucleotide addition rate (green), Pause 1 (dark blue) and Pause 2 (cyan) exit rates for the conditions described in (B). **(D)** Probabilities to enter Pause 1 (dark blue), Pause 2 (cyan) and the backtrack (red) states for the conditions described in (B) and Fig. 3D. **(E)** Dwell time distributions of SARS-CoV-2 polymerase activity traces for 500 µM all other NTPs without (gray) or with 300 µM T1106-TP (red). The corresponding solid lines are the fit to the pause-stochastic model. **(F)** Nucleotide addition rate (green), Pause 1 (dark blue) and Pause 2 (cyan) exit rates for the conditions described in (E). **(G)** Probabilities to enter Pause 1 (dark blue), Pause 2 (cyan) and the backtrack (red) states for the conditions described in (E) and Fig. 4D. The error bars in (B, E) represent one standard deviation extracted from 1000 bootstraps. The error bars in (C, E, F, G) are one standard deviation extracted from 100 bootstraps.

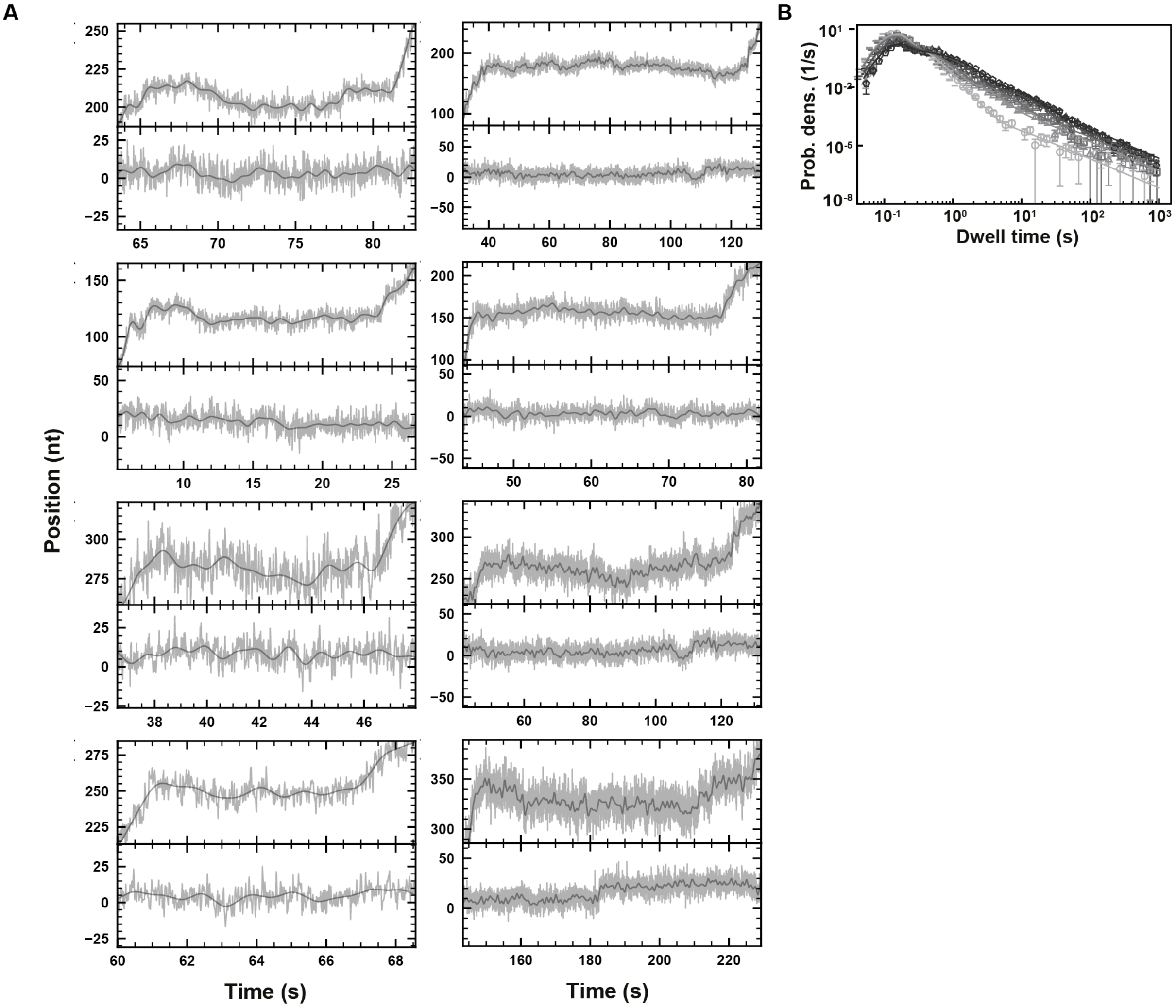


**Figure S5: SARS-CoV-2 polymerase elongation traces in presence of RDV-TP at 25°C. (A)** Examples of deep SARS-CoV-2 backtracks induced by RDV-TP incorporation (top) and traces showing no polymerase activity (bottom). Traces acquired using an ultra-stable magnetic tweezers as described in Ref. (*12*) at 35 pN, 58 Hz acquisition frequency (grey), low-pass filtered at 1 Hz (dark grey), and using a reaction buffer containing 10 µM RDV-TP, 50 µM ATP and 500 µM all other NTPs. **(B)** Dwell time distributions of SARS-CoV-2 polymerase activity traces acquired in the presence of 500 µM NTP at 25°C, either without (circles), or with 20 µM (triangles), 50 µM (squares), 100 µM (diamonds), and 300 µM (pentagons) RDV-TP. The color code from light to dark gray further highlights the increasing concentration of RDV-TP. The solid lines represent the fit of the pause-stochastic model. The error bars represent one standard deviation extracted from 1000 bootstraps.

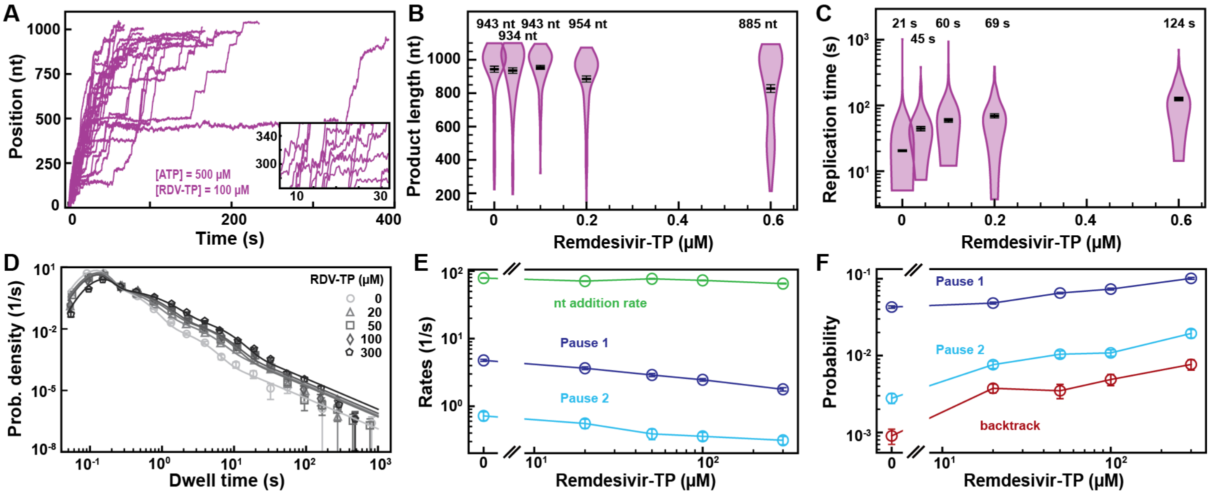


**Figure S6: SARS-CoV-1 polymerase activity traces kinetics in presence of RDV-TP. (A)** SARS-CoV-1 replication traces for 500 µM NTPs and 100 µM RDV-TP. **(B)** SARS-CoV-1 polymerase product length for the 1043 nt long template using 500 µM NTPs, as a function of [RDV-TP]/[ATP]. The mean values are indicated above the violin plots, and represented by horizontal black thick lines flanked by one standard deviation error bars extracted from 1000 bootstraps. **(C)** Replication time for the reaction conditions described in (B). The median values are indicated above the violin plots, and represented by horizontal black thick lines flanked by one standard deviation error bars extracted from 1000 bootstraps. **(D)** Dwell time distributions of SARS-CoV-1 polymerase activity traces acquired in the presence of 500 µM NTP, either without (circles), or with 20 µM (triangles), 50 µM (squares), 100 µM (diamonds), and 300 µM (pentagons) RDV-TP. The color code from light to dark gray further highlights the increasing concentration of RDV-TP. The solid lines represent the fit of the pause-stochastic model. **(E)** Nucleotide addition rate (green), Pause 1 (dark blue) and Pause 2 (cyan) exit rates from the fit of the dwell time distributions in (B). **(F)** Probabilities to enter Pause 1 (dark blue), Pause 2 (cyan) and the backtrack (red) states for the conditions described in (B). The error bars in (D) represent one standard deviation extracted from 1000 bootstraps. The error bars in (E, F) are one standard deviation extracted from 100 bootstrap.

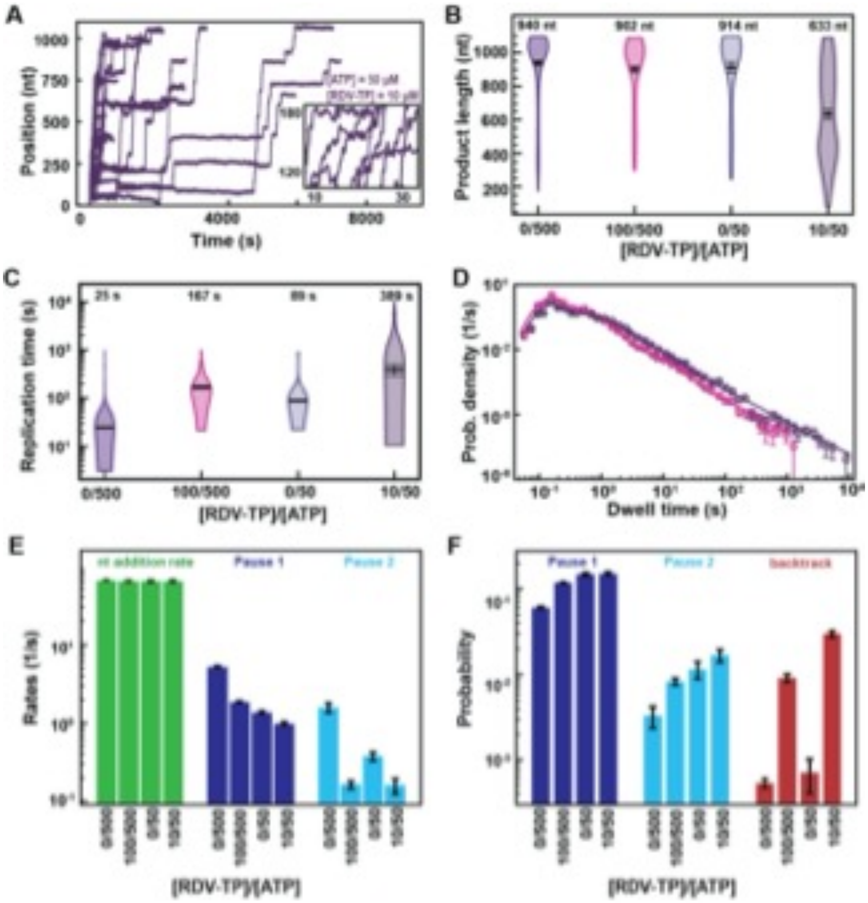


**Figure S7: Lower ATP concentration at constant RDV-TP:ATP stoichiometry increases the effects of RDV-TP on SARS-CoV-2 polymerase elongation kinetics. (A)** SARS-CoV-2 polymerase activity traces for 10 µM RDV-TP with 50 µM ATP and 500 µM of the other NTPs. **(B)** Product length of SARS-CoV-2 polymerase at RDV-TP:ATP stoichiometry of 0/500, 100/500, 0/50 and 10/50. The mean values are indicated above the violin plots, and represented by horizontal black thick lines flanked by one standard deviation error bars extracted from 1000 bootstraps. **(C)** The replication times for the conditions described in (B). The median values are indicated above the violin plots, and represented by horizontal black thick lines flanked by one standard deviation extracted from 1000 bootstraps. **(D)** Dwell time distributions of SARS-CoV-2 polymerase activity traces for either 50 µM ATP, 500 µM all other NTPs and 10 µM RDV-TP (purple), or at 500 µM all NTPs and 100 µM RDV-TP (pink). The corresponding solid lines are the fit to the pause-stochastic model. **(E)** Nucleotide addition rate (green), Pause 1 (dark blue) and Pause 2 (cyan) exit rates for the conditions described in (B). **(F)** Probabilities to enter Pause 1 (dark blue), Pause 2 (cyan) and the backtrack (red) states for the conditions described in (B). The error bars in (D) represent one standard deviation extracted from 1000 bootstraps. The error bars in (E, F) are one standard deviation extracted from 100 bootstraps.

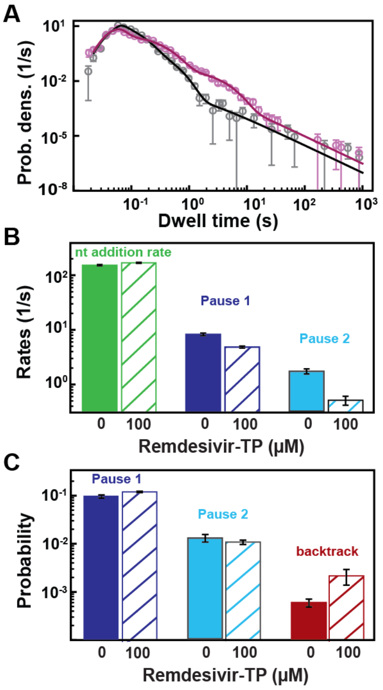


**Figure S8: SARS-CoV-2 polymerase activity traces kinetics in presence of RDV-TP at 25°C and 37°C. (A)** Dwell time distributions of SARS-CoV-2 replication activity acquired in the presence of 500 µM NTP at 37°C without (gray) and with 100 µM RDV-TP (pink). The solid lines represent the fit of the pause-stochastic model. **(B)** Nucleotide addition rate (green), Pause 1 (dark blue) and Pause 2 (cyan) exit rates from the fit of the dwell time distributions in (A). **(C)** Probabilities to enter Pause 1 (dark blue), Pause 2 (cyan) and the backtrack (red) states for the conditions described in (A). The error bars in (A) represent one standard deviation extracted from 1000 bootstraps. The error bars in (B,C) are one standard deviation extracted from 100 bootstraps.

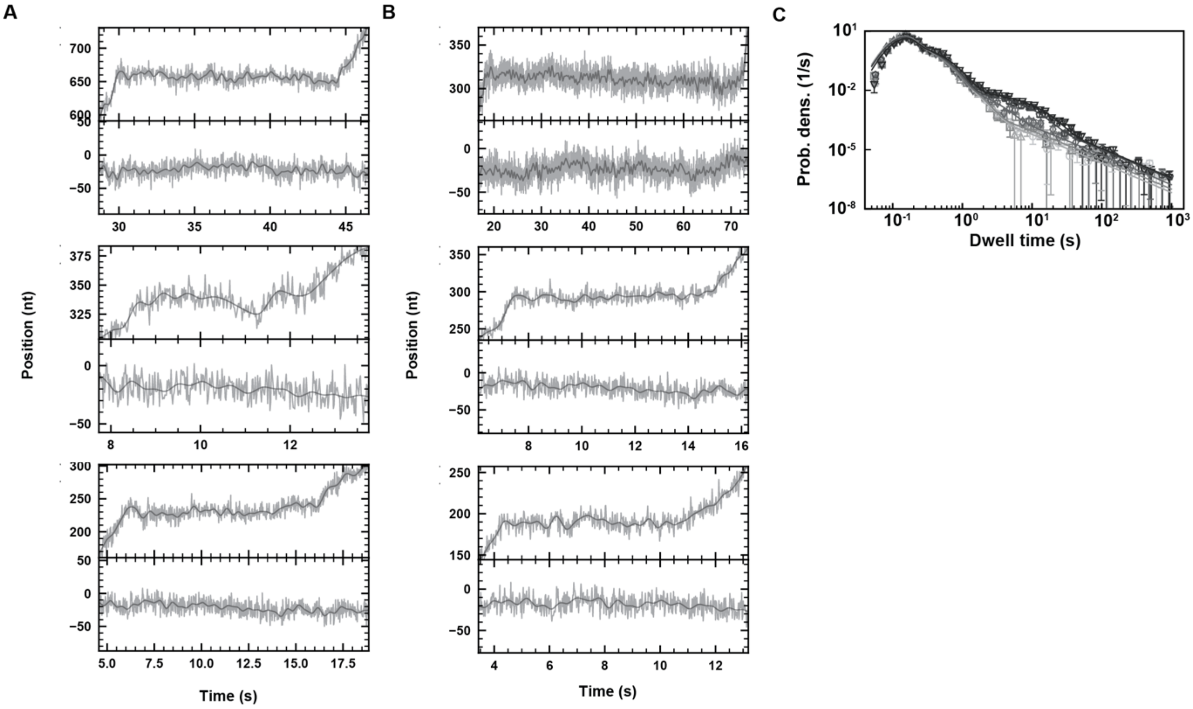


**Figure S9: SARS-CoV-2 polymerase elongation in presence of T-1106-TP.** Using an ultra-stable magnetic tweezers configuration, we monitored pauses in SARS-CoV-2 polymerase activity traces at 58 Hz camera acquisition frequency (grey; 1 Hz low-pass filtered: dark grey), applying 35 pN force, and in the presence of 500 µM T-1106-TP and 500 µM all NTPs. Top: zoom in the polymerase activity traces; bottom inactive tether followed simultaneously. **(A)** Some pauses demonstrate shallow backtracks, **(B)** while in other pause cases, polymerase backtrack is difficult to confirm given the spatiotemporal resolution of the assay. **(C)** Dwell time distributions of SARS-CoV-2 polymerase activity traces acquired in the presence of 500 µM NTP, either without (circles), or with 20 µM (triangles), 50 µM (squares), 100 µM (diamonds), 300 µM (pentagons), and 500 µM (upside down triangle) T-1106-TP. The color code from light to dark gray further highlights the increasing concentration of T-1106-TP. The solid lines represent the fit of the pause-stochastic model. The error bars represent one standard deviation extracted from 1000 bootstraps.

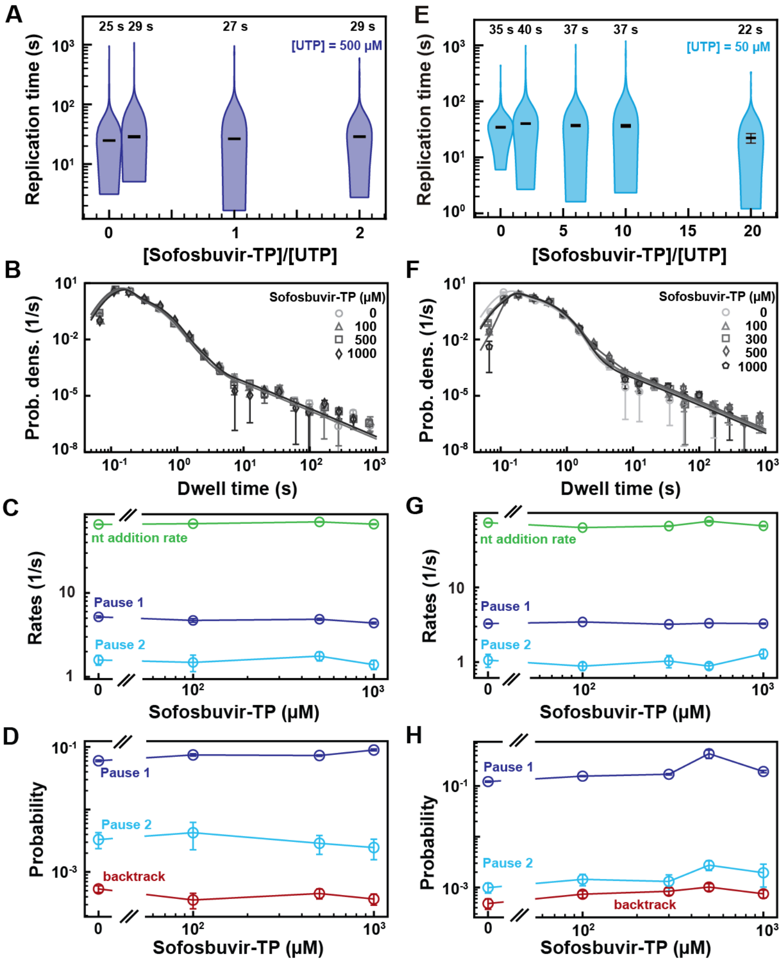


**Figure S10: SARS-CoV-2 polymerase activity traces kinetics in presence of Sofosbuvir-TP. (A, E)** SARS-CoV-2 replication time for the 1043 nt long template using the indicated concentration of UTP, 500 µM of other NTPs as a function of the stoichiometry of [Sofosbuvir-TP]/[UTP]. The median values are indicated above the violin plots, and represented by horizontal black thick lines flanked by one standard deviation extracted from 1000 bootstraps. **(B)** Dwell time distributions of SARS-CoV-2 polymerase activity traces acquired in the presence of 500 µM NTP either without (circles) or with 0.1 mM (triangles), 0.5 mM (squares), and 1 mM (diamonds) Sofosbuvir-TP. The color code from light to dark gray further highlights the increasing concentration of Sofosbuvir-TP. The solid lines represent the fit to the pause-stochastic model. **(C)** Nucleotide addition rate (green), Pause 1 (dark blue) and Pause 2 (cyan) exit rates for the conditions described in (B). **(D)** Probabilities to enter Pause 1 (dark blue), Pause 2 (cyan) and the backtrack (red) states for the conditions described in (B). **(F)** Dwell time distributions of SARS-CoV-2 polymerase activity traces acquired in the presence of 50 µM UTP, 500 µM of other NTPs, and either without (circles), or with 0.1 mM (triangles), 0.3 mM (squares), 0.5 mM (diamonds) and 1 mM (pentagons) Sofosbuvir-TP. The color code from light to dark gray further highlights the increasing concentration of Sofosbuvir-TP. The solid lines represent the fit to the pause-stochastic model. **(G)** Nucleotide addition rate (green), Pause 1 (dark blue) and Pause 2 (cyan) exit rates for the conditions described in (F). **(H)** Probabilities to enter Pause 1 (dark blue), Pause 2 (cyan) and the backtrack (red) states for the conditions described in (F). The error bars in (B and F) represent one standard deviation extracted from 1000 bootstraps. The error bars in (C, D, G, H) are one standard deviation extracted from 100 bootstraps.

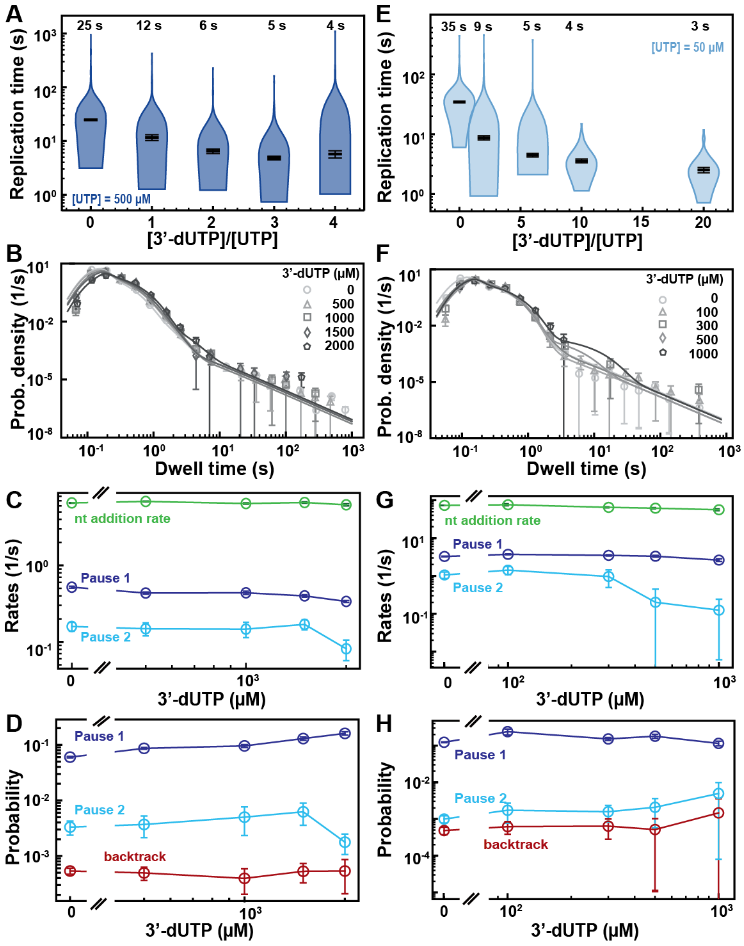


**Figure S11: SARS-CoV-2 polymerase activity traces kinetics in presence 3'-dUTP. (A, E)** SARS-CoV-2 replication time for the 1043 nt long template using the indicated concentration of UTP, 500 µM of other NTPs as a function of the stoichiometry of [3'-dUTP]/[UTP]. The median values are indicated above the violin plots, and represented by horizontal black thick lines flanked by one standard deviation extracted from 1000 bootstraps. **(B)** Dwell time distributions of SARS-CoV-2 polymerase activity traces acquired in the presence of 500 µM NTP either without (circles) or with 0.5 mM (triangles), 1 mM (squares), 1.5 mM (diamonds), and 2 mM (pentagon) 3'-dUTP. The color code from light to dark gray further highlights the increasing concentration of 3'-dUTP. The solid lines represent the fit to the pause-stochastic model. **(C)** Nucleotide addition rate (green), Pause 1 (dark blue) and Pause 2 (cyan) exit rates for the conditions described in (B). **(D)** Probabilities to enter Pause 1 (dark blue), Pause 2 (cyan) and the backtrack (red) states for the conditions described in (B). **(F)** Dwell time distributions of SARS-CoV-2 polymerase activity traces acquired in the presence of 50 µM UTP, 500 µM of other NTPs, and either without (circles), or with 0.1 mM (triangles), 0.3 mM (squares), 0.5 mM (diamonds) and 1 mM (pentagons) 3'-dUTP. The color code from light to dark gray further highlights the increasing concentration of 3'-dUTP. The solid lines represent the fit to the pause-stochastic model. **(G)** Nucleotide addition rate (green), Pause 1 (dark blue) and Pause 2 (cyan) exit rates for the conditions described in (F). **(H)** Probabilities to enter Pause 1 (dark blue), Pause 2 (cyan) and the backtrack (red) states for the conditions described in (F). The error bars in (B and F) represent one standard deviation extracted from 1000 bootstraps. The error bars in (C, D, G, H) are one standard deviation extracted from 100 bootstraps.

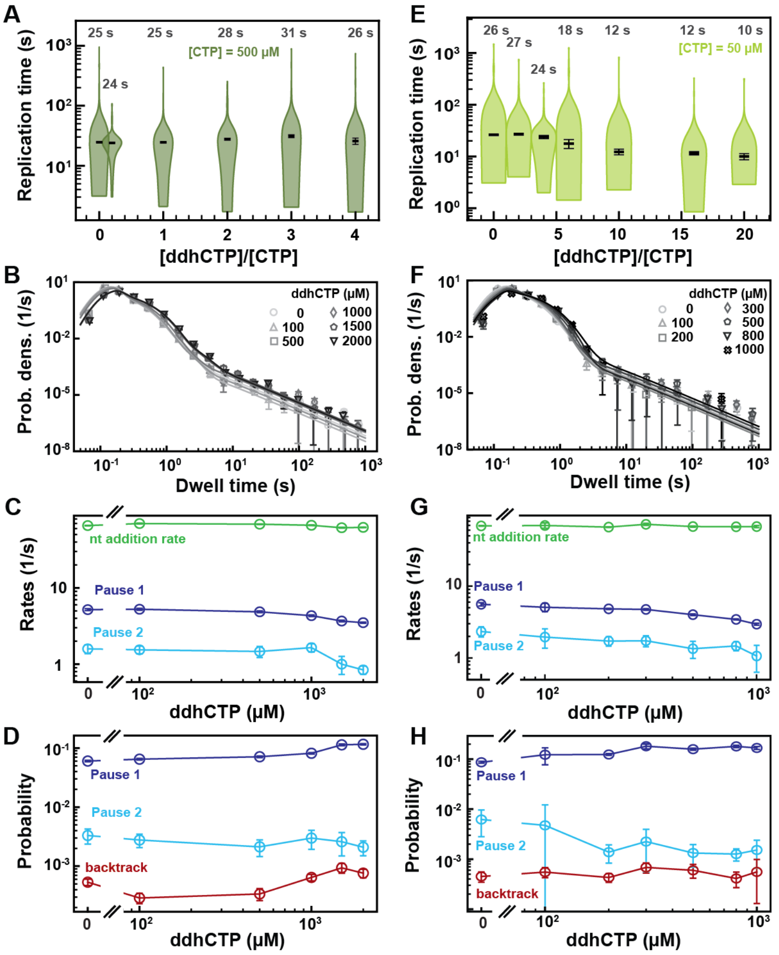


**Figure S12: SARS-CoV-2 polymerase activity traces in presence of ddhCTP. (A, E)** SARS-CoV-2 replication time for the 1043 nt long template using the indicated concentration of CTP, 500 µM of other NTPs, and the indicated stoichiometry of [ddhCTP]/[CTP]. The median values are indicated above the violin plots, and represented by horizontal black thick lines flanked by one standard deviation extracted from 1000 bootstraps. **(B)** Dwell time distributions of SARS-CoV-2 polymerase activity traces acquired in the presence of 500 µM NTP either without (circles) or with 0.1 mM (triangles), 0.5 mM (squares), and 1 mM (diamonds) ddhCTP. The color code from light to dark gray further highlights the increasing concentration of ddhCTP. The solid lines represent the fit to the pause-stochastic model. **(C)** Nucleotide addition rate (green), Pause 1 (dark blue) and Pause 2 (cyan) exit rates for the conditions described in (B). **(D)** Probabilities to enter Pause 1 (dark blue), Pause 2 (cyan) and the backtrack (red) states for the conditions described in (B). **(F)** Dwell time distributions of SARS-CoV-2 polymerase activity traces acquired in the presence of 50 µM CTP, 500 µM of other NTPs, and either without (circles), or with 0.1 mM (triangles), 0.2 mM (squares), 0.3 mM (diamonds), 0.5 mM (pentagons), 0.8 mM (upside down triangle) and 1 mM (x) ddhCTP. The color code from light to dark gray further highlights the increasing concentration of ddhCTP. The solid lines represent the fit to the pause-stochastic model. **(G)** Nucleotide addition rate (green), Pause 1 (dark blue) and Pause 2 (cyan) exit rates for the conditions described in (F). **(H)** Probabilities to enter Pause 1 (dark blue), Pause 2 (cyan) and the backtrack (red) states for the conditions described in (F). The error bars in (B and F) represent one standard deviation extracted from 1000 bootstraps. The error bars in (C, D, G, H) are one standard deviation extracted from 100 bootstraps.

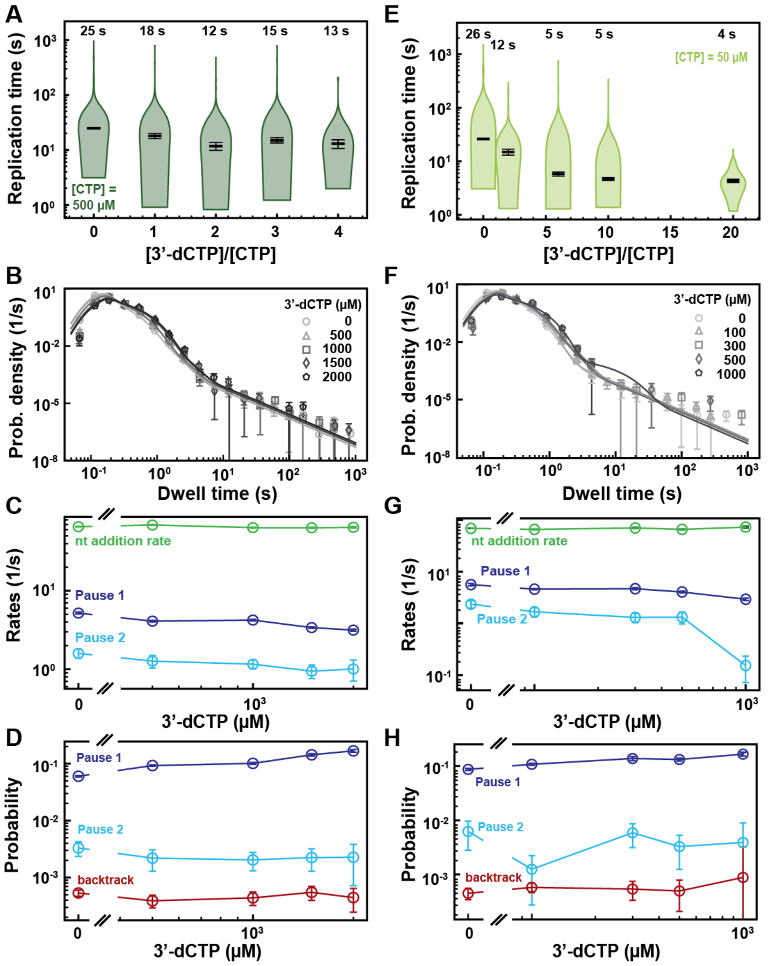


**Figure S13: SARS-CoV-2 polymerase activity traces kinetics in presence 3'-dCTP. (A, E)** SARS-CoV-2 replication time for the 1043 nt long template using the indicated concentration of CTP, 500 µM of other NTPs as a function of the stoichiometry of [3'-dCTP]/[CTP]. The median values are indicated above the violin plots, and represented by horizontal black thick lines flanked by one standard deviation extracted from 1000 bootstraps. **(B)** Dwell time distributions of SARS-CoV-2 polymerase activity traces acquired in the presence of 500 µM NTP either without (circles) or with 0.5 mM (triangles), 1 mM (squares), 1.5 mM (diamonds), and 2 mM (pentagon) 3'-dCTP. The color code from light to dark gray further highlights the increasing concentration of 3'-dCTP. The solid lines represent the fit to the pause-stochastic model. **(C)** Nucleotide addition rate (green), Pause 1 (dark blue) and Pause 2 (cyan) exit rates for the conditions described in (B). **(D)** Probabilities to enter Pause 1 (dark blue), Pause 2 (cyan) and the backtrack (red) states for the conditions described in (B). **(F)** Dwell time distributions of SARS-CoV-2 polymerase activity traces acquired in the presence of 50 µM CTP, 500 µM of other NTPs, and either without (circles), or with 0.1 mM (triangles), 0.3 mM (squares), 0.5 mM (diamonds) and 1 mM (pentagons) 3'-dCTP. The color code from light to dark gray further highlights the increasing concentration of 3'-dCTP. The solid lines represent the fit to the pause-stochastic model. **(G)** Nucleotide addition rate (green), Pause 1 (dark blue) and Pause 2 (cyan) exit rates for the conditions described in (F). **(H)** Probabilities to enter Pause 1 (dark blue), Pause 2 (cyan) and the backtrack (red) states for the conditions described in (F). The error bars in (B and F) represent one standard deviation extracted from 1000 bootstraps. The error bars in (C, D, G, H) are one standard deviation extracted from 100 bootstraps.

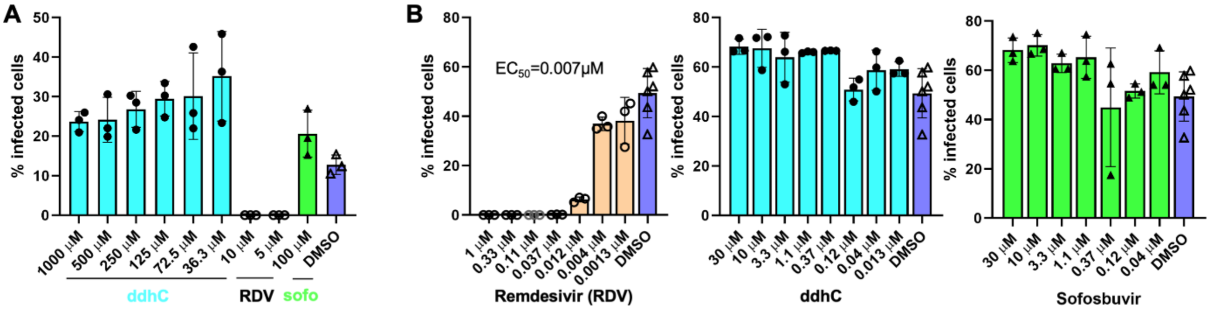


**Figure S14: ddhC does not inhibit SARS-CoV-2 replication in huh7-hACE2 cells.** Huh7-hACE2 cells in 96-wells were incubated with the indicated concentrations of the tested compounds for 1.5 hours before SARS-CoV-2 (USA-WA1/2020 isolate) was added at MOI of 0.05. At ~24 hpi, the percentage of infected cells was assessed by immunofluorescence assay using a rabbit monoclonal antibody against the SARS-CoV-2 N protein. “sofo” is Sofosbuvir. Results in (A) and (B) represent two separate experiments.

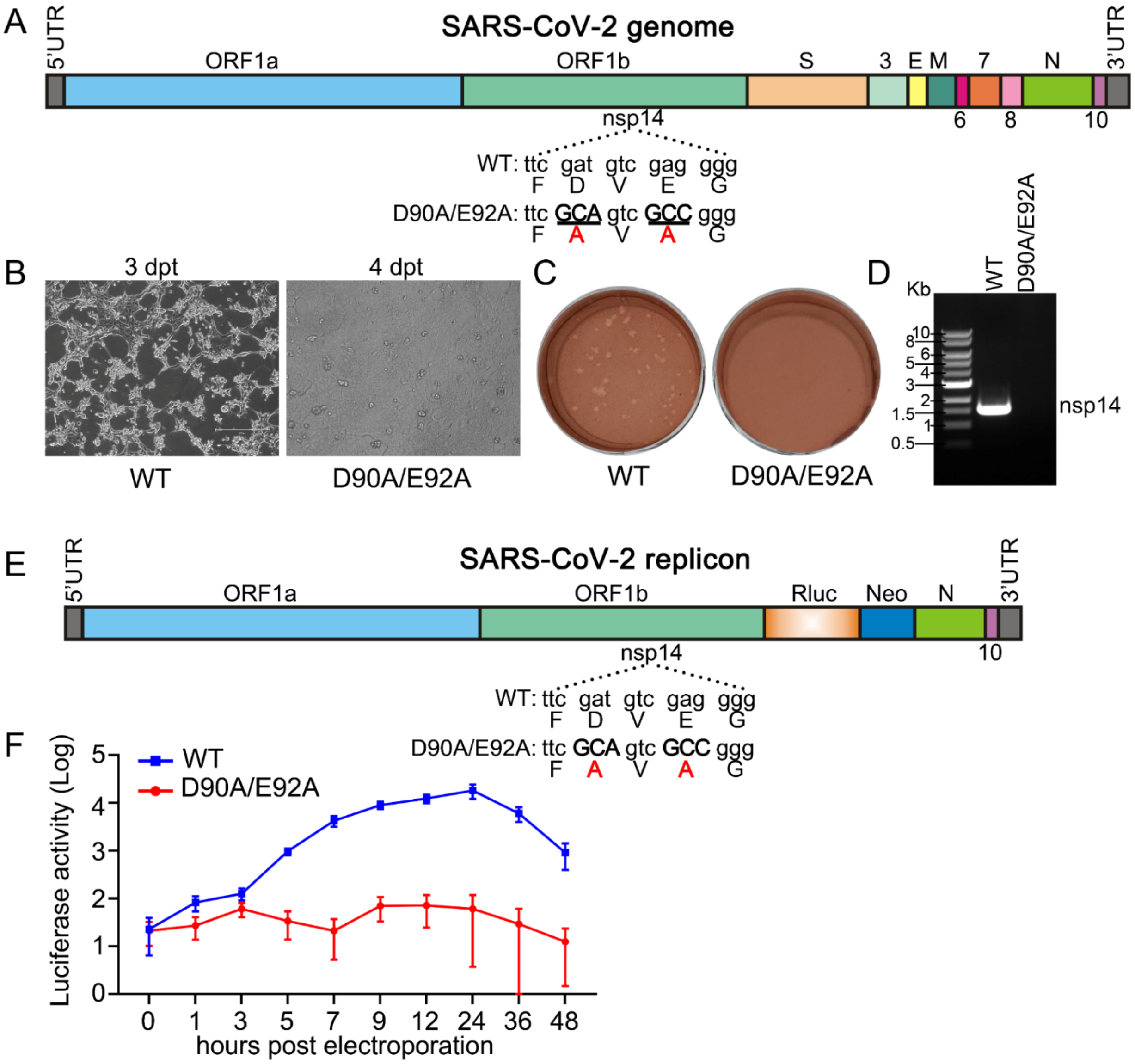


**Figure S15: SARS-CoV-2 nsp14 exoribonuclease knockout is not replicative. (A)** SARS-CoV-2 genome. The SARS-CoV-2 nsp14 exoribonuclease nucleotides and amino acid mutations (D90A/E92A) are indicated. **(B)** Phase-contrast images of electroporated cells. Vero E6 cells were electroporated with SARS-CoV-2 WT or nsp14 exoribonuclease knockout mutant RNA. **(C)** Plaque morphology of SARS-CoV-2 WT and nsp14 exoribonuclease knockout mutant viruses. Supernatants were harvested on day 3 post-electroporation (WT) and day 4 post-electroporation (mutant) from **(B)**. Plaque assay was performed in Vero E6 cells and staining with neutral red solution after 48 h infection. **(D)** RT-PCR analysis. Extracellular RNA from **(B)** were harvested on day 3 (WT) and day 4 (mutant). The nsp14 region of SARS-CoV-2 was amplified by RT-PCR to confirm viral production. **(E)** Transient replicon of SARS-CoV-2. A *Renilla* luciferase gene is inserted as a reporter and the nsp14 exoribonuclease knockout is shown as indicated. **(F)** Replicon luciferase assay. Huh-7 cells were electroporated with WT or mutant replicon RNA, cells were harvested and assayed for luciferase activities at indicated timepoints.

### Table S1

| nucleotide analog | NTP conc. (µM) | nucleotide analog conc. (µM) | T (°C) | Force (pN) | product length: # traces | product length: (mean ± std) nt | product length: (γ ± std) nt | total replication time (median ± std) s | dwell time distribution: # traces | # dwell times | (nucleotide addition rate ± std) 1/s | (pause 1 exit rate ± std) 1/s | (pause 2 exit rate ± std) 1/s | pause 1 probability ± std | pause 2 probability ± std | backtrack probability ± std | Figures |
|---|---|---|---|---|---|---|---|---|---|---|---|---|---|---|---|---|---|
| 3'-dATP | 500 A/C/G/UTP | 0 | 25 | 35 | 269 | 940 ± 13 | 780 ± 64 | 25 ± 1 | 343 | 29458 | 65.6 ± 0.5 | 5.19 ± 0.14 | 1.59 ± 0.21 | 0.060 ± 0.002 | 0.0033 ± 0.0009 | 0.0005 ± 0.0001 | 1BCE-H, 2A-C, 3B-E, 4B-F, 5CD, 6CD, S1D, S3A-C, S5, S6B-F, S8BCEF, S9, S10A-D, S11A-D, S12A-D, S13A-D |
| | | 500 | | | 97 | 802 ± 30 | | 23 ± 1 | 98 | 7238 | 66.2 ± 1.6 | 5.74 ± 0.76 | 2.66 ± 0.51 | 0.067 ± 0.005 | 0.0129 ± 9.0084 | 0.0004 ± 0.0001 | 2ABC, S3A-C |
| | | 1000 | | | 133 | 701 ± 25 | | 27 ± 2 | 134 | 8478 | 63.5 ± 0.8 | 3.85 ± 0.10 | 1.60 ± 0.20 | 0.010 ± 0.004 | 0.0015 ± 0.0008 | 0.0004 ± 0.0001 | 2BC, S3A-C |
| | | 1500 | | | 146 | 643 ± 26 | | 29 ± 3 | 154 | 9131 | 56.1 ± 1.4 | 3.42 ± 0.06 | 1.06 ± 0.15 | 0.155 ± 0.009 | 0.0021 ± 0.0007 | 0.0006 ± 0.0001 | 2BC, S3A-C |
| | | 2000 | | | 98 | 566 ± 33 | | 24 ± 4 | 98 | 5155 | 61.3 ± 1.4 | 3.08 ± 0.08 | 1.09 ± 0.15 | 0.177 ± 0.012 | 0.0036 ± 0.0015 | 0.0004 ± 0.0002 | 2BC, S3A-C |
| | 500 C/G/UTP, 50 ATP | 0 | 25 | 35 | 61 | 914 ± 29 | 777 ± 50 | 89 ± 3 | 61 | 4808 | 64.0 ± 1.2 | 1.38 ± 0.05 | 0.37 ± 0.05 | 0.149 ± 0.005 | 0.0115 ± 0.0026 | 0.0007 ± 0.0003 | 2DEF, S3D-F, S7BCEF |
| | | 100 | | | 60 | 698 ± 38 | | 83 ± 10 | 62 | 3855 | 62.6 ± 1.1 | 1.26 ± 0.05 | 0.33 ± 0.04 | 0.152 ± 0.005 | 0.0109 ± 0.0024 | 0.0014 ± 0.0005 | 2EF, S3D-F |
| | | 300 | | | 69 | 383 ± 28 | | 28 ± 4 | 69 | 2241 | 62.8 ± 2.1 | 1.43 ± 0.11 | 0.50 ± 0.07 | 0.140 ± 0.009 | 0.0225 ± 0.0067 | 0.0012 ± 0.0006 | 2EF, S3D-F |
| | | 500 | | | 60 | 299 ± 27 | | 23 ± 3 | 60 | 1442 | 61.9 ± 1.9 | 1.36 ± 0.18 | 0.45 ± 0.16 | 0.154 ± 0.016 | 0.0229 ± 0.0130 | 0.0025 ± 0.0042 | 2EF, S3D-F |
| | | 1000 | | | 51 | 195 ± 16 | | 19 ± 3 | 51 | 737 | 77.5 ± 5.7 | 1.22 ± 0.08 | 0.28 ± 0.06 | 0.213 ± 0.025 | 0.0126 ± 0.0039 | 0.0026 ± 0.0025 | 2EF, S3D-F |
| | 500 C/G/UTP, 50 ATP | 300 | 25 | 25 | 44 | 341 ± 33 | - | 28 ± 6 | - | | | | | | | | S4A |
| 3'-dCTP | 500 A/C/G/UTP | 0 | 25 | 35 | 269 | 940 ± 13 | 338 ± 18 | 25 ± 1 | 343 | 29458 | 65.6 ± 0.5 | 5.19 ± 0.14 | 1.59 ± 0.21 | 0.060 ± 0.002 | 0.0033 ± 0.0009 | 0.0005 ± 0.0001 | 1BCE-H, 2A-C, 3B-E, 4B-F, 5CD, 6CD, S1D, S3A-C, S5, S6B-F, S8BCEF, S9, S10A-D, S11A-D, S12A-D, S13A-D |
| | | 500 | | | 148 | 604 ± 27 | | 18 ± 2 | 162 | 9383 | 69.0 ± 0.7 | 4.10 ± 0.09 | 1.27 ± 0.23 | 0.093 ± 0.003 | 0.0022 ± 0.0009 | 0.0004 ± 0.0001 | 6BD, S12A-D |
| | | 1000 | | | 159 | 482 ± 27 | | 12 ± 2 | 162 | 7242 | 63.8 ± 0.8 | 4.23 ± 0.10 | 1.16 ± 0.23 | 0.101 ± 0.004 | 0.0020 ± 0.0007 | 0.0004 ± 0.0001 | 6D, S13A-D |
| | | 1500 | | | 139 | 440 ± 23 | | 15 ± 2 | 157 | 6063 | 63.5 ± 1.0 | 3.39 ± 0.08 | 0.94 ± 0.19 | 0.143 ± 0.007 | 0.0022 ± 0.0009 | 0.0006 ± 0.0002 | 6D, S13A-D |
| | | 2000 | | | 101 | 352 ± 24 | | 13 ± 2 | 110 | 3609 | 64.4 ± 1.2 | 3.15 ± 0.09 | 1.01 ± 0.30 | 0.169 ± 0.009 | 0.0023 ± 0.0016 | 0.0004 ± 0.0002 | 6D, S13A-D |
| | 500 A/G/UTP, 50 CTP | 0 | 25 | 35 | 79 | 966 ± 12 | 457 ± 21 | 26 ± 1 | 82 | 7593 | 68.7 ± 0.7 | 6.68 ± 0.30 | 2.33 ± 0.38 | 0.087 ± 0.004 | 0.0062 ± 0.0034 | 0.0005 ± 0.0001 | 6GH, S13E-H, S13E-H |
| | | 100 | | | 171 | 565 ± 25 | | 15 ± 2 | 190 | 9727 | 65.6 ± 0.8 | 4.56 ± 0.10 | 1.66 ± 0.34 | 0.107 ± 0.004 | 0.0013 ± 0.0010 | 0.0006 ± 0.0001 | 6H, S13E-H |
| | | 300 | | | 129 | 263 ± 16 | | 6 ±1 | 133 | 3078 | 69.9 ± 2.0 | 4.65 ± 0.22 | 1.30 ± 0.27 | 0.137 ± 0.012 | 0.0059 ± 0.0027 | 0.0005 ± 0.0002 | 6H, S13E-H |
| | | 500 | | | 108 | 178 ± 11 | | 5 ± 0 | 112 | 1638 | 65.6 ± 1.3 | 4.02 ± 0.15 | 1.31 ± 0.34 | 0.132 ± 0.008 | 0.0033 ± 0.0020 | 0.0005 ± 0.0003 | 6FH, S13E-H |
| | | 1000 | | | 68 | 131 ± 8 | | 4 ± 0 | 71 | 688 | 72.7 ± 2.9 | 2.90 ± 0.15 | 0.15 ± 0.08 | 0.165 ± 0.013 | 0.0039 ± 0.0050 | 0.0009 ± 0.0024 | 6H, S13E-H |
| 3'-dUTP | 500 A/C/G/UTP | 0 | 25 | 35 | 269 | 940 ± 13 | 151 ± 6 | 25 ± 1 | 343 | 29458 | 65.6 ± 0.5 | 5.19 ± 0.14 | 1.59 ± 0.21 | 0.060 ± 0.002 | 0.0033 ± 0.0009 | 0.0005 ± 0.0001 | 1BCE-H, 2A-C, 3B-E, 4B-F, 5CD, 6CD, S1D, S3A-C, S5, S6B-F, S8BCEF, S9, S10A-D, S11A-D, S12A-D, S13A-D |
| | | 500 | | | 130 | 445 ± 26 | | 12 ± 1 | 151 | 6188 | 68.5 ± 0.9 | 4.35 ± 0.14 | 1.48 ± 0.29 | 0.086 ± 0.003 | 0.0037 ± 0.0015 | 0.0005 ± 0.0001 | 5AD, S11A-D |
| | | 1000 | | | 141 | 278 ± 17 | | 6 ± 1 | 155 | 3933 | 64.4 ± 1.2 | 4.38 ± 0.21 | 1.47 ± 0.33 | 0.096 ± 0.006 | 0.0050 ± 0.0026 | 0.0004 ± 0.0002 | 5D, S11A-D |
| | | 1500 | | | 148 | 210 ± 14 | | 5 ± 0 | 160 | 2859 | 66.4 ± 1.3 | 3.99 ± 0.14 | 1.69 ± 0.25 | 0.130 ± 0.009 | 0.0062 ± 0.0027 | 0.0005 ± 0.0002 | 5D, S11A-D |
| | | 2000 | | | 100 | 197 ± 14 | | 6 ± 1 | 114 | 1821 | 61.9 ± 2.0 | 3.38 ± 0.09 | 0.81 ± 0.24 | 0.161 ± 0.011 | 0.0018 ± 0.0007 | 0.0005 ± 0.0003 | 5D, S11A-D |
| | 500 A/C/GTP, 50 UTP | 0 | 25 | 35 | 98 | 910 ± 22 | 241 ± 9 | 34 ± 1 | 117 | 9742 | 73.5 ± 0.9 | 3.27 ± 0.05 | 1.06 ± 0.21 | 0.122 ± 0.003 | 0.0010 ±0.0002 | 0.0005 ± 0.0001 | 5GH, S9E-H, S11E-H |
| | | 100 | | | 135 | 315 ± 21 | | 9 ± 1 | 143 | 3849 | 76.3 ± 4.6 | 3.72 ± 0.09 | 1.43 ± 0.35 | 0.238 ± 0.044 | 0.0017 ± 0.0010 | 0.0006 ± 0.0002 | 5H, S11E-H |
| | | 300 | | | 92 | 146 ± 7 | | 4 ± 0 | 94 | 1094 | 65.1 ± 1.8 | 3.50 ± 0.14 | 0.97 ± 0.47 | 0.151 ± 0.014 | 0.0015 ± 0.0007 | 0.0006 ± 0.0004 | 5H, S11E-H |
| | | 500 | | | 74 | 108 ± 7 | | 4 ± 0 | 76 | 627 | 61.9 ± 2.3 | 3.31 ± 0.17 | 0.20 ± 0.24 | 0.179 ± 0.022 | 0.0021 ± 0.0015 | 0.0005 ± 0.0005 | 5FH, S11E-H |
| | | 1000 | | | 57 | 67 ± 3 | | 3 ± 0 | 60 | 260 | 56.3 ± 3.2 | 2.62 ± 0.21 | 0.12 ± 0.12 | 0.114 ± 0.017 | 0.0049 ± 0.0049 | 0.0015 ± 0.0021 | 5H, S11E-H |
| ddhCTP | 500 A/C/G/UTP | 0 | 25 | 35 | 269 | 940 ± 13 | 1221 ± 130 | 25 ± 1 | 343 | 29458 | 65.6 ± 0.5 | 5.19 ± 0.14 | 1.59 ± 0.21 | 0.060 ± 0.002 | 0.0033 ± 0.0009 | 0.0005 ± 0.0001 | 1BCE-H, 2A-C, 3B-E, 4B-F, 5CD, 6CD, S1D, S3A-C, S5, S6B-F, S8BCEF, S9, S10A-D, S11A-D, S12A-D, S13A-D |
| | | 100 | | | 172 | 934 ± 16 | | 24 ± 1 | 213 | 18749 | 69.8 ± 0.7 | 5.25 ± 0.15 | 1.54 ± 0.18 | 0.065 ± 0.002 | 0.0028 ± 0.0007 | 0.0003 ± 0.0001 | 6C, S12A-D |
| | | 500 | | | 125 | 894 ± 22 | | 25 ± 1 | 145 | 11877 | 68.4 ± 0.6 | 4.86 ± 0.13 | 1.47 ± 0.24 | 0.071 ± 0.002 | 0.0021 ± 0.0003 | 0.0003 ± 0.0001 | 6AC, S12A-D |
| | | 1000 | | | 146 | 789 ± 25 | | 28 ± 1 | 152 | 11213 | 66.2 ± 0.6 | 4.33 ± 0.10 | 1.64 ± 0.20 | 0.082 ± 0.002 | 0.0030 ± 0.0011 | 0.0006 ± 0.0001 | 6C, S12A-D |
| | | 1500 | | | 162 | 742 ± 26 | | 31 ± 2 | 168 | 11796 | 61.5 ± 0.8 | 3.68 ±0.09 | 1.00 ± 0.27 | 0.113 ± 0.004 | 0.0026 ± 0.0011 | 0.0009 ± 0.0002 | 6C, S12A-D |
| | | 2000 | | | 187 | 622 ± 24 | | 26 ± 3 | 190 | 11003 | 62.1 ± 0.6 | 3.50 ± 0.06 | 0.84 ± 0.11 | 0.116 ± 0.003 | 0.0021 ± 0.0006 | 0.0008 ± 0.0001 | 6C, S12A-D |
| | 500 A/G/UTP, 50 CTP | 0 | 25 | 35 | 79 | 966 ± 19 | 1360 ± 71 | 26 ± 1 | 82 | 7593 | 68.7 ± 0.7 | 5.58 ± 0.30 | 2.33 ± 0.38 | 0.087 ± 0.004 | 0.0062 ± 0.0034 | 0.0005 ± 0.0001 | 6GH, S12E-H, S13E-H |
| | | 100 | | | 69 | 815 ± 32 | | 27 ± 1 | 77 | 5708 | 69.7 ± 5.9 | 5.07 ± 0.68 | 1.95 ± 0.58 | 0.122 ± 0.046 | 0.0047 ± 0.0074 | 0.0006 ± 0.0001 | 6G, S12E-H |
| | | 200 | | | 152 | 700 ± 26 | | 24 ± 2 | 167 | 11151 | 66.1 ± 0.6 | 4.82 ± 0.08 | 1.72 ± 0.26 | 0.124 ± 0.005 | 0.0014 ± 0.0005 | 0.0004 ± 0.0001 | 6EG, S12E-H |
| | | 300 | | | 86 | 599 ± 31 | | 18 ± 3 | 94 | 5111 | 72.3 ± 2.5 | 4.73 ± 0.12 | 1.74 ± 0.30 | 0.181 ± 0.021 | 0.0022 ± 0.0017 | 0.0007 ± 0.0002 | 6G, S12E-H |
| | | 500 | | | 79 | 433 ± 29 | | 12 ± 2 | 83 | 3139 | 67.3 ± 1.1 | 4.00 ± 0.10 | 1.34 ± 0.36 | 0.158 ± 0.008 | 0.0013 ± 0.0006 | 0.0006 ± 0.0002 | 6G, S12E-H |
| | | 800 | | | 157 | 362 ± 18 | | 12 ± 1 | 157 | 4938 | 66.8 ± 1.3 | 3.44 ± 0.06 | 1.47 ± 0.21 | 0.180 ± 0.010 | 0.0013 ± 0.0003 | 0.0004 ± 0.0001 | 6G, S12E-H |
| | | 1000 | | | 59 | 284 ± 23 | | 10 ± 1 | 61 | 1480 | 67.3 ± 1.1 | 2.95 ± 0.10 | 1.06 ± 0.43 | 0.167 ± 0.011 | 0.0015 ± 0.0009 | 0.0006 ± 0.0004 | 6G, S12E-H |
| | 500 A/G/UTP, 50 CTP | 300 | 25 | 25 | 92 | 499 ± 31 | - | 13 ± 2 | - | | | | | | | | S4A |

#### Table S2:

| nucleotide analog | NTP conc. (µM) | nucleotide analog conc. (µM) | T (°C) | Force (pN) | product length # traces | product length (mean ± std) nt | product length (γ ± std) nt | total replication time (median ± std) s | dwell time distribution # traces | # dwell times | (nucleotide addition rate ± std) 1/s | (pause 1 exit rate ± std) 1/s | (pause 2 exit rate ± std) 1/s | pause 1 probability ± std | pause 2 probability ± std | backtrack probability ± std | Figures |
|---|---|---|---|---|---|---|---|---|---|---|---|---|---|---|---|---|---|
| Remdesivir-TP SARS-CoV-2 | 500 A/C/G/UTP | 0 | 25 | 35 | 269 | 940 ± 13 | - | 25 ± 1 | 343 | 29458 | 65.6 ± 0.5 | 5.19 ± 0.14 | 1.59 ± 0.21 | 0.060 ± 0.002 | 0.0033 ± 0.0009 | 0.0005 ± 0.0001 | 1BCE-H, 2A-C, 3B-E, 4B-F, 5CD, 6CD, S1D, S3A-C, S5, S6B-F, S8BCEF, S9, S10A-D, S11A-D, S12A-D, S13A-D |
| | | 20 | | | 114 | 976 ± 14 | | 53 ± 2 | 148 | 12409 | 67.9 ± 0.8 | 4.13 ± 0.16 | 0.62 ± 0.11 | 0.074 ± 0.002 | 0.0062 ± 0.0009 | 0.0030 ± 0.0004 | 3BCEF, S5B |
| | | 50 | | | 87 | 947 ± 23 | | 72 ± 6 | 148 | 11710 | 67.3 ± 0.7 | 3.39 ± 0.12 | 0.41 ± 0.07 | 0.080 ± 0.002 | 0.0071 ± 0.0007 | 0.0052 ± 0.0006 | 3BCEF, S5B |
| | | 100 | | | 120 | 902 ± 18 | | 167 ± 9 | 138 | 10350 | 64.1 ± 0.7 | 1.87 ± 0.04 | 0.16 ± 0.02 | 0.120 ± 0.003 | 0.0083 ± 0.0007 | 0.0091 ± 0.0010 | 3A-F, S5, S8B-F |
| | | 300 | | | 106 | 894 ± 20 | | 315 ± 14 | 161 | 11544 | 61.4 ± 0.8 | 1.38 ± 0.04 | 0.16 ± 0.03 | 0.140 ± 0.003 | 0.0121 ± 0.0001 | 0.0142 ± 0.0015 | 3BCEF, S5B |
| | 500 A/C/G/UTP | 0 | 37 | 35 | 78 | 909 ± 31 | - | 12 ± 1 | 87 | 7101 | 169.0 ± 3.8 | 15.60 ± 0.71 | 4.37 ± 0.39 | 0.132 ± 0.010 | 0.0163 ± 0.0030 | 0.0006 ± 0.0001 | 1BE-H, S1D, S7A-C |
| | | 100 | | | 52 | 768 ± 34 | | 33 ± 4 | 57 | 4057 | 169.2 ± 3.3 | 4.85 ± 0.19 | 0.51 ± 0.10 | 0.119 ± 0.004 | 0.0108 ± 0.0011 | 0.0022 ± 0.0007 | S7A-C |
| | 500 A/C/G/UTP | 0 | 25 | 25 | 93 | 924 ± 12 | - | 21 ± 1 | 191 | 16262 | 77.7 ± 0.5 | 5.47 ± 0.13 | 1.74 ± 0.32 | 0.062 ± 0.002 | 0.0018 ± 0.0007 | 0.0008 ± 0.0001 | S4A-G |
| | | 100 | | | 113 | 892 ± 15 | | 91 ± 6 | 131 | 10194 | 72.1 ± 0.5 | 2.19 ± 0.05 | 0.37 ± 0.04 | 0.091 ± 0.002 | 0.0086 ± 0.0010 | 0.0049 ± 0.0005 | S4B-D |
| | 500 C/G/UTP, 50 ATP | 0 | 25 | 35 | 61 | 914 ± 29 | - | 89 ± 3 | 61 | 4808 | 64.0 ± 1.2 | 1.38 ± 0.05 | 0.37 ± 0.05 | 0.149 ± 0.005 | 0.0115 ± 0.0026 | 0.0007 ± 0.0003 | 2DEF, S3D-F, S8BCEF |
| | | 10 | | | 90 | 633 ± 30 | | 389 ± 106 | 96 | 5244 | 64.1 ± 1.5 | 0.98 ± 0.05 | 0.16 ± 0.04 | 0.151 ± 0.006 | 0.0167 ± 0.0028 | 0.0299 ± 0.0023 | S8A-F |
| Sofosbuvir-TP | 500 A/C/G/UTP | 0 | 25 | 35 | 269 | 940 ± 13 | - | 25 ± 1 | 343 | 29458 | 65.6 ± 0.5 | 5.19 ± 0.14 | 1.59 ± 0.21 | 0.060 ± 0.002 | 0.0033 ± 0.0009 | 0.0005 ± 0.0001 | 1BCE-H, 2A-C, 3B-E, 4B-F, 5CD, 6CD, S1D, S3A-C, S5, S6B-F, S8BCEF, S9, S10A-D, S11A-D, S12A-D, S13A-D |
| | | 100 | | | 108 | 967 ± 19 | | 29 ± 1 | 126 | 11185 | 67.0 ± 0.7 | 4.71 ± 0.23 | 1.49 ± 0.33 | 0.075 ± 0.003 | 0.0043 ± 0.0020 | 0.0004 ± 0.0001 | 5C, S10A-D |
| | | 500 | | | 128 | 966 ± 17 | | 27 ± 1 | 158 | 14364 | 70.4 ± 0.5 | 4.86 ± 0.13 | 1.77 ± 0.07 | 0.073 ± 0.002 | 0.0029 ± 0.0010 | 0.0005 ± 0.0001 | 5C, S10A-D |
| | | 1000 | | | 131 | 934 ± 19 | | 29 ± 1 | 166 | 14879 | 66.2 ± 0.7 | 4.38 ± 0.10 | 1.40 ± 0.18 | 0.089 ± 0.003 | 0.0025 ± 0.0009 | 0.0004 ± 0.0001 | 5AC, S10A-D |
| | 500 A/C/GTP, 50 UTP | 0 | 25 | 35 | 98 | 910 ± 21 | 3908 ± 467 | 35 ± 1 | 117 | 9742 | 73.5 ± 0.9 | 3.27 ± 0.05 | 1.06 ± 0.21 | 0.123 ± 0.003 | 0.0010 ± 0.0002 | 0.0005 ± 0.0001 | 5GH, S10E-H, S11E-H |
| | | 100 | | | 148 | 873 ± 24 | | 40 ± 1 | 167 | 13448 | 63.1 ± 0.5 | 3.44 ± 0.04 | 0.88 ± 0.11 | 0.157 ± 0.004 | 0.0014 ± 0.0004 | 0.0007 ± 0.0001 | 5G, S10E-H |
| | | 300 | | | 122 | 734 ± 30 | | 37 ± 2 | 143 | 9757 | 66.1 ± 0.9 | 3.21 ± 0.04 | 1.03 ± 0.19 | 0.171 ± 0.005 | 0.0013 ± 0.0005 | 0.0008 ± 0.0001 | 5G, S10E-H |
| | | 500 | | | 167 | 706 ± 27 | | 37 ± 2 | 188 | 12238 | 76.7 ± 1.9 | 3.31 ± 0.04 | 0.88 ± 0.08 | 0.435 ± 0.085 | 0.0028 ± 0.0006 | 0.0010 ± 0.0002 | 5EG, S10E-H |
| | | 1000 | | | 127 | 563 ± 32 | | 22 ± 4 | 138 | 7170 | 66.7 ± 1.2 | 3.27 ± 0.06 | 1.30 ± 0.19 | 0.193 ± 0.011 | 0.0020 ± 0.0009 | 0.0008 ± 0.0001 | 5G, S10E-H |
| | 500 A/C/GTP, 50 UTP | 300 | 25 | 25 | 69 | 709 ± 34 | - | 30 ± 1 | - | | | | | | | | S4A |
| T-1106-TP | 500 A/C/G/UTP | 0 | 25 | 35 | 269 | 940 ± 13 | - | 25 ± 1 | 343 | 29458 | 65.6 ± 0.5 | 5.19 ± 0.14 | 1.59 ± 0.21 | 0.060 ± 0.002 | 0.0033 ± 0.0009 | 0.0005 ± 0.0001 | 1BCE-H, 2A-C, 3B-E, 4B-F, 5CD, 6CD, S1D, S3A-C, S5, S6B-F, S8BCEF, S9, S10A-D, S11A-D, S12A-D, S13A-D |
| | | 20 | | | 131 | 945 ± 20 | | 28 ± 1 | 173 | 14922 | 69.7 ± 0.5 | 5.19 ± 0.14 | 1.52 ± 0.14 | 0.073 ± 0.002 | 0.0033 ± 0.0007 | 0.0005 ± 0.0001 | 4BCEF, S9C |
| | | 50 | | | 137 | 929 ± 16 | | 27 ± 1 | 174 | 15041 | 67.8 ± 0.7 | 5.12 ± 0.11 | 1.89 ± 0.19 | 0.071 ± 0.004 | 0.0066 ± 0.0015 | 0.0007 ± 0.0001 | 4BCEF, S9C |
| | | 100 | | | 131 | 943 ± 18 | | 30 ± 1 | 166 | 14159 | 67.7 ± 0.5 | 6.25 ± 0.29 | 0.77 ± 0.13 | 0.071 ± 0.002 | 0.0025 ± 0.0004 | 0.0011 ± 0.0002 | 4BCEF, S9C |
| | | 300 | | | 145 | 931 ± 17 | | 52 ± 4 | 160 | 12987 | 68.9 ± 0.6 | 4.98 ± 0.10 | 0.22 ± 0.03 | 0.079 ± 0.002 | 0.0035 ± 0.0003 | 0.0021 ± 0.0004 | 4BCEF, S9C |
| | | 500 | | | 144 | 946 ± 16 | | 65 ± 4 | 171 | 14626 | 64.2 ± 0.4 | 3.83 ± 0.07 | 0.16 ± 0.01 | 0.079 ± 0.002 | 0.0044 ± 0.0003 | 0.0023 ± 0.0004 | 4A-F, S9C |
| | 500 A/C/G/UTP | 0 | 25 | 25 | 93 | 924 ± 12 | - | 21 ± 0 | 191 | 16262 | 77.7 ± 0.5 | 5.47 ± 0.13 | 1.74 ± 0.32 | 0.062 ± 0.002 | 0.0018 ± 0.0007 | 0.0008 ± 0.0001 | S4E-G |
| | | 300 | | | 84 | 916 ± 16 | | 44 ± 2 | 142 | 11673 | 70.2 ± 0.4 | 4.08 ± 0.10 | 0.20 ± 0.04 | 0.062 ± 0.002 | 0.0027 ± 0.0004 | 0.0019 ± 0.0007 | S4E-G |
| Remdesivir-TP SARS-CoV-1 | 500 A/C/G/UTP | 0 | 25 | 35 | 93 | 943 ± 20 | - | 21 ± 1 | 107 | 9525 | 78.7 ± 0.5 | 4.76 ± 0.16 | 0.72 ± 0.11 | 0.043 ± 0.001 | 0.0028 ± 0.0004 | 0.0009 ± 0.0002 | 1BCE-H, 2A-C, 3B-E, 4B-F, 5CD, 6CD, S1D, S3A-C, S5, S6B-F, S8BCEF, S9, S10A-D, S11A-D, S12A-D, S13A-D |
| | | 20 | | | 117 | 934 ± 17 | | 45 ± 3 | 148 | 13193 | 71.1 ± 0.5 | 3.64 ± 0.17 | 0.56 ± 0.09 | 0.048 ± 0.001 | 0.0076 ± 0.0009 | 0.0038 ± 0.0005 | S6B-F |
| | | 50 | | | 124 | 954 ± 12 | | 60 ± 4 | 137 | 12300 | 76.7 ± 0.7 | 2.90 ± 0.15 | 0.39 ± 0.07 | 0.065 ± 0.001 | 0.0104 ± 0.0011 | 0.0035 ± 0.0007 | S6B-F |
| | | 100 | | | 98 | 885 ± 19 | | 69 ± 5 | 111 | 8951 | 72.8 ± 0.7 | 2.46 ± 0.10 | 0.36 ± 0.05 | 0.073 ± 0.002 | 0.0109 ± 0.0010 | 0.0049 ± 0.0008 | S6A-F |
| | | 300 | | | 116 | 827 ± 18 | | 124 ± 8 | 135 | 10422 | 65.2 ± 0.8 | 1.78 ± 0.10 | 0.31 ± 0.04 | 0.101 ± 0.003 | 0.0194 ± 0.0024 | 0.0077 ± 0.0012 | S6B-F |
